# Spatio-temporal X-linked gene reactivation and site-specific retention of epigenetic silencing in the mouse germline

**DOI:** 10.1101/2023.04.25.532252

**Authors:** Clara Roidor, Laurène Syx, Emmanuelle Beyne, Dina Zielinski, Aurélie Teissandier, Caroline Lee, Marius Walter, Nicolas Servant, Karim Chebli, Déborah Bourc’his, M. Azim Surani, Maud Borensztein

## Abstract

Random X-chromosome inactivation (XCI) is a hallmark of female mammalian somatic cells. This epigenetic mechanism, mediated by the long non-coding RNA *Xist*, occurs in the epiblast and is stably maintained to ensure proper dosage compensation of X-linked genes during life. However, this silencing is lost during primordial germ cell (PGC) development. Using a combination of single-cell allele-specific RNA sequencing and low-input chromatin profiling in developing *in vivo* PGC, we provide unprecedented detailed maps of gene reactivation. We demonstrated that PGC still carry a fully silent X chromosome on embryonic day (E) 9.5, despite the loss of *Xist* expression. X-linked genes are then gradually reactivated outside the *Xist* first-bound regions. At E12.5, a significant part of the inactive X chromosome (Xi) still resists reactivation, carrying an epigenetic memory of its silencing. Late-reactivated genes are enriched in repressive chromatin marks, including DNA methylation and H3K27me3 marks. Our results define the timing of reactivation of the silent X chromosome a key event in female PGC reprogramming with direct implications for reproduction.

## Introduction

In mammals, while proper commitment and homeostasis of somatic lineages are central to individual survival, correct establishment of the germline is crucial for functional gamete and species survival. In mice, specification of germ cell lineage is initiated during embryonic post-implantation development at the onset of gastrulation. Approximately 30-40 PGC become specified and are found at the base of the allantois bud at E7.25 ^1, 2^. After E8.5, PGC undergo migration and proliferation, and reach the genital ridges between E10 and E11. Throughout this period, germ cell proliferation and colonization continue until the PGC enters meiotic prophase at E13.5 in the female embryos, a process that occurs only after birth in males ^3^. Establishment of the germline determines the production of oocytes and spermatozoa, and therefore their competency to accomplish fertilization and transmit genetic and epigenetic information to the next generation ^4^. PGC differentiation is accompanied by repression of the somatic program, expression of germline-specific genes, and extensive genome-wide epigenetic reprogramming including DNA demethylation, loss of genomic imprints, and redistribution of histone marks ^1, 5^. Epigenetic reprogramming occurs when PGCs proliferate and migrate to colonize the future gonads. During the upstream development of gonadal sex determination, PGC epigenetic reprogramming displays striking differences between XX females and XY males, with reactivation of the inactive X chromosome ^6–9^. This leads to an excess of X-linked gene products, with the X:Autosome ratio exceeding 1 in female PGCs compared to males ^10, 11^. Excess X-linked genes could promote sexual dimorphism and meiosis progression through the direct or indirect involvement of some X-linked genes in the process of sex-specific gonadal formation and Xi reactivation itself ^8, 12^.

Differences in sex chromosome content between males and females lead to gene dosage imbalance. This is compensated in mammals by transcriptional silencing of one of the two X chromosomes in female somatic cells ^13, 14^. This epigenetic mechanism, called X-chromosome inactivation, represents an important paradigm for chromosome-wide epigenetic regulation. Long non-coding RNA *Xist* plays a crucial role in the initiation of XCI ^15–17^. Its absence in early female embryos leads to lethality owing to both impaired dosage compensation and extra-embryonic tissue development ^16, 17^. Once expressed from the future Xi, *Xist* coats in cis the most accessible regions, in 3D spatial proximity, the *Xist* ‘entry sites’, before spreading along chromosome ^18–20^. Transposons, particularly LINE1 elements, have been proposed to facilitate this heterochromatinization process ^21, 22^. *Xist* then triggers the recruitment of specific factors involved in gene silencing, which in turn induces the removal of active chromatin marks and recruitment of repressive histone marks, such as H3K27me3, H3K9me2, and H2AK119ub ^23^. Finally, DNA methylation is recruited to the promoters of inactivated genes to further lock the silent state of these genes and maintain them over hundreds of cell divisions ^24^. Studies on the kinetics of XCI have shown that different genes followed different speeds of silencing in the pre-implantation ^16^ and post-implantation mouse embryos ^25^, and during *in vitro* differentiation of mouse embryonic stem cells (mES) ^26, 27^, except for a small subset of genes that resist silencing (escapee, ∼7 % in the mouse). Early silent genes seem to be more prone to lie inside *Xist* ‘entry sites’ and close to the X-inactivation centre (Xic) ^16^ from which *Xist* is transcribed, in gene-rich regions, pre-bound by Polycomb ^23^ and close to LINE1 elements ^27^.

Although XCI has been extensively studied over the past 60 years, much less is known about how Xi reactivation occurs and whether it mirrors XCI key events. Upon development, Xi reactivation occurs after imprinted XCI in the inner cell mass (ICM) of the blastocyst ^28, 29^ (a rodent-specific event) and in PGC ^6, 7^ after random XCI. It is also observed *in vitro* during female induced pluripotent stem cell derivation (iPSC) ^30, 31^. Based on a combination of immunofluorescence, RNA-FISH, and RT-PCR in the germline, early PGC carry an inactive X chromosome enriched in H3K27me3^7, 9, 32^, with silent X-linked genes ^6^. Reactivation is initiated by the transcriptional extinction of *Xist* RNA (from E7.5, prior to and upon PGC migration), loss of repressive H3K27me3 chromatin marks (from E9.5, during PGC migration and proliferation), and re-expression of silenced X-linked genes (from E10.5, when PGC start to colonize the future gonads) ^6, 7, 32, 33^. *Xist* repression has been linked to pluripotency factors such as Nanog and Prdm14 in pluripotent mES ^34^, ICM ^29, 35^ and germline ^9^.

However, studies based on single-cell RNA sequencing (scRNA-seq) in post-implantation embryos have shown progressive random XCI depending on lineage ^25, 36^. Some epiblast cells can still carry two active X chromosomes at the time of PGC specification. A recent study on Xi reactivation in *in vitro* PGC-like cells suggested that Xi could only be moderately silenced in early PGC before full reactivation ^8^. This study was based on *in vitro* XCI in Epiblast-like cells differentiated from mES followed by Xi reactivation in induced PGC-like cells.

Despite this knowledge, the events underlying X-linked gene reactivation in the germline remain unclear, particularly at the level of the entire chromosome. The extent of XCI in early *in vivo* PGC and the kinetics of gene reactivation are still unknown. To address these questions, we explored the precise kinetics of X-linked gene expression and Xi chromatin change. We combined interspecific mouse crosses and scRNA-seq, DNA methylation assay using Whole Genome Bisulfite (WGBS), and H3K27me3 histone mark profiling using low-input allele-specific CUT&RUN. We investigated the transcriptional changes from E8.5 to E12.5 PGC. We showed that X-linked genes are sequentially activated, as previously described for the ICM of the blastocyst^29^ but with different dynamics and requirements. In PGCs, we observed reactivation dependency on *Xist* entry sites, DNA methylation levels, H3K27me3 enrichment, and genomic location. This study provides important insights into the transcriptional, allelic, and chromatin dynamics of Xi reactivation in germline cells. Our novel results emphasize the importance of studying epigenetic reprogramming of the inactive X chromosome in the unique context of the germline, which is the most relevant for developmental syndromes and human reproductive medicine.

## Results

### Transcriptional analysis of *in vivo* PGC and soma cells by scRNA-seq

To address the X-chromosome reactivation kinetics in PGC, at the chromosome-wide scale, we produced high-quality, high-coverage scRNA-seq for female and male PGC, as well as surrounding female somatic cells, as a control for the maintenance of X-chromosome inactivation. F1 interspecific embryos were obtained by mating *Mus Musculus domesticus* (129) inbred and *Mus Musculus castaneus* (Cast) inbred mice, and collected every day between E8.5 and E12.5 (**Figure 1A**). These sub-species have evolved for more than 3 million years and carry an important number of well-characterized single nucleotide polymorphisms (SNP) (mean of ∼1 SNP / 650 bp on the X chromosome between 129 and Cast, Mouse Genomes project) ^37, 38^. To visualize PGC in our embryos, we utilized Green Fluorescent Protein (GFP) transgene, in a 129 genetic background under the control of *Stella* or *Oct4* promoter. *Stella* gene (also known as *Dppa3*) is expressed early during PGC development, at the time of emergence ^39^. We used it for E8.5 embryos. After E8.5, we took advantage of the Oct4-GFP transgenic mice ^40^. *Oct4* also known as

**Figure 1:**
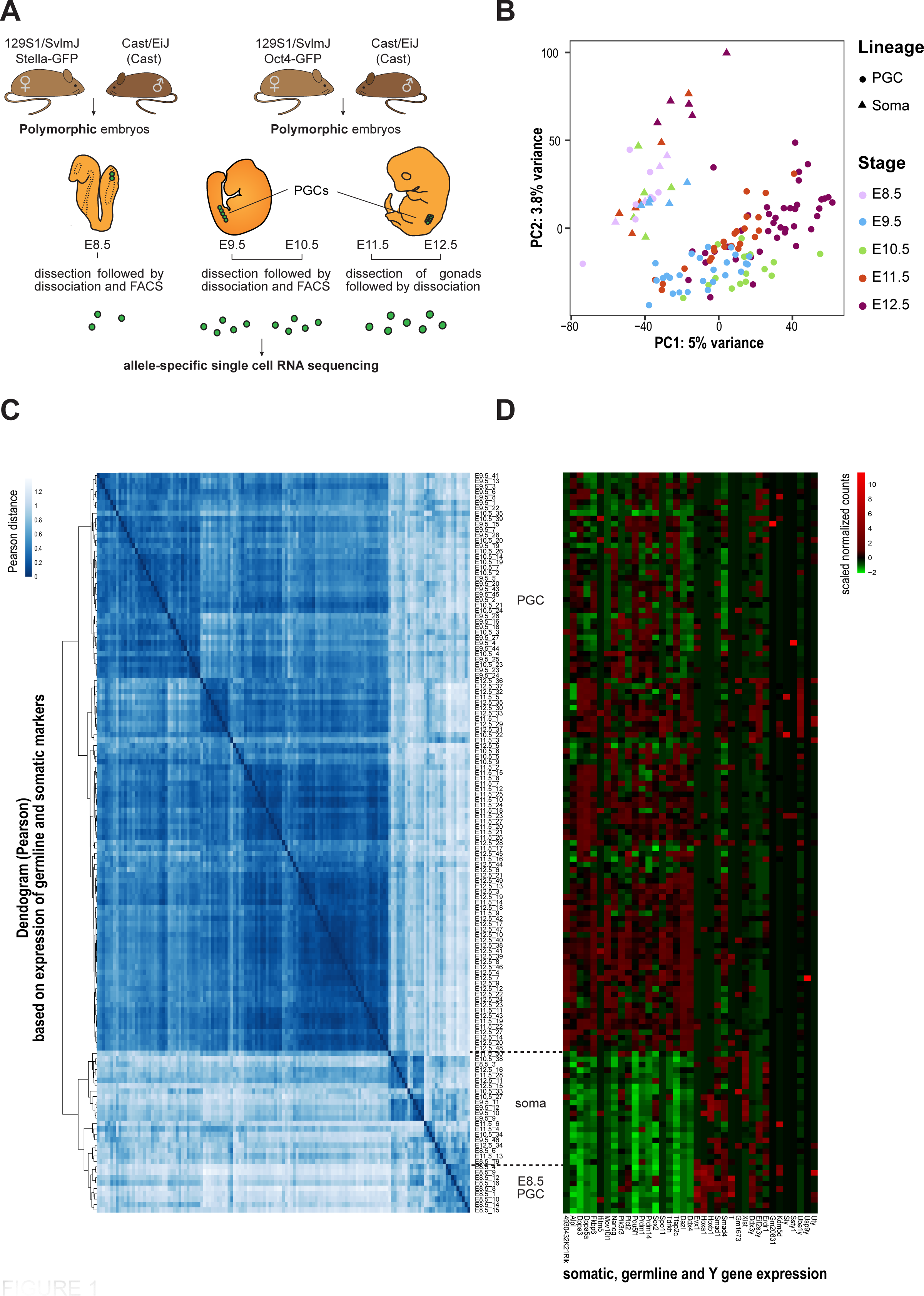
Single-cell RNA sequencing of polymorphic primordial germ cells during embryonic development. (**A**) Schematic illustration of single-cell transcriptomic experiments including mouse breeding (129 × cast), harvested embryonic stages, and single-cell collection. At E8.5, PGC were sorted according to the expression of *Stella*-GFP. *Stella* gene, also known as *Dppa3*, is expressed in early PGC. From E9.5, *Oct4-GFP* (GOF-△PE-18 line)^40^ marker was used to differentiate between PGC and soma. PGC, primordial germ cells; FACS, fluorescence-activated cell sorting. **(B)** Principal component analysis (PCA) of single PGC and soma based on the 1 000 most variable genes in the transcriptomic datasets. The different stages are denoted by different colours. The rounds represent PGC, and triangles represent GFP-negative somatic cells in the proximity of the PGC. The number of cells analysed per stage and further details of the scRNA-seq samples are shown in Supplementary Table 1. **(C)** Hierarchical clustering and Pearson’s distance of scRNA-seq samples based on germline, soma, and sex-specific gene expression variation using Pearson’s correlation. Cells were clustered first by lineage (PGC and soma), then by stage (E8.5–E12.5), and then by sex for the E11.5 and E12.5 stages. n = 137 single-cell samples. **(D)** Expression levels of 26 known genes expressed in developing PGC or soma, and sex-specific genes (*Xist* and Y-linked genes) in the 137 single-cell samples were used to classify cells according to their lineage, as shown. The cells were ordered according to hierarchical clustering in **C**. PGC primordial germ cells.

*Pou5f1*, encodes a pluripotent transcription factor present in PGC, but not in surrounding somatic cells. From E8.5 to E10.5, PGC cells were collected with the assistance of fluorescent active cell sorting based on GFP expression. From E11.5, soma and PGC cells were collected after gonad dissection, based on their size, and confirmed based on the presence of GFP under a microscope. Each single cell was then manually picked, and poly adenylated mRNA was amplified ^42^. We produced high-quality scRNA-seq libraries from 154 samples. Since we were interested in allelic expression, we decided to use a scRNA-seq method ^41^ allowing high-depth of high-throughput sequencing for each single cell (**Supplemental Table 1**). Only single cells that passed the quality controls (see Methods) were used for downstream analysis (n=137 libraries). We used Principal Component Analysis (PCA) to associate single cells based on their lineage (PGC versus soma, **Figure 1B**). The cells were first associated based on their lineage, followed by their embryonic stage. Next, we performed a correlation analysis based on the expression status of pluripotency, soma, and Y-linked genes (**Figures 1B and C**; genes listed in **Figure 1C**). We classified the cells according to their developmental stage and pluripotency/soma factor status. This clearly supports a strong repression of the somatic program in PGC and confirmed the expression of well-known factors in PGC such as Stella, Blimp1, Oct4, Nanog, and Dazl ^1, 43^. The weight of these known factors in segregating the soma and PGC lineages is shown in **Extended Figure 1A**. The length of the arrows is proportional to the implication of the factor in PCA association. Without surprise, markers of soma point towards somatic cells and germline towards PGC. With a closer look it is important to note that late PGC genes preferentially point towards E12.5 PGC. We then asked the best 30 predictor genes of PCA clustering (**Extended Figure 1B)**. Some well-known factors were found to push towards PGC fate, such as Oct4, Tfap2c, and DND1 ^1, 44^. However, other genes with interesting roles have also been identified, such as Zing Finger Protein Zfp985. The ZFP family is important for protecting DNA methylation at imprinting loci and transposons ^45^. Recently, Zfp982 has been associated with the stemness state of mES through its potential control of Nanog and Stella ^46^. Epithelial splicing regulatory protein 1, Esrp1, induces oocyte defects and female infertility if deleted from E15 ^47^. Our dataset could reveal novel PGC markers and are important candidates for PGC biology.

We then studied the sex of our single-cell samples to focus on sex specificities, X-chromosome reactivation, and transcriptional changes during female PGC development. After clustering PGC and soma cells based on the expression of well-known markers and confirmation by clustering analysis (**Figures 1C and D**), we sexed all single cells based on *Xist* (in XX females) and Y-linked gene (in XY male) expression, allowing attribution to each single cell towards a lineage and sex (**Figure 2A**, **Supplemental Table 1**). Sex was also confirmed by the absence of SNPs from the X chromosomes in XY cells. By serendipity, an XO female embryo was found at E12.5, and was used in the analysis (n= 3 PGC and 1 soma cells sequenced). We first studied the differentially expressed genes (DEG) in migratory and colonizing female PGC by comparing E9.5 versus E10.5, E11.5 versus E10.5 and E12.5 versus E11.5 (**Extended Figures 1C, 1D and 1E**). Strikingly, very few DEG were found in migratory PGC (E10.5 versus E9.5). This is in accordance with a previously published large scRNA-seq dataset of mouse male and female PGC ^43^. From E11.5, there was a strong increase in DEG in female PGC, with expected upregulation of PGC genes such as *Dazl* and *Stella*/*Dppa3*. In accordance with previous reports, there was significant gene repression and upregulation at E12.5, compared to E11.5 ^43, 48^. Most DEG from the X-chromosome were upregulated, which is consistent with X-chromosome reactivation occurring in female PGC (**Extended Figure 1E**).

**Figure 2:**
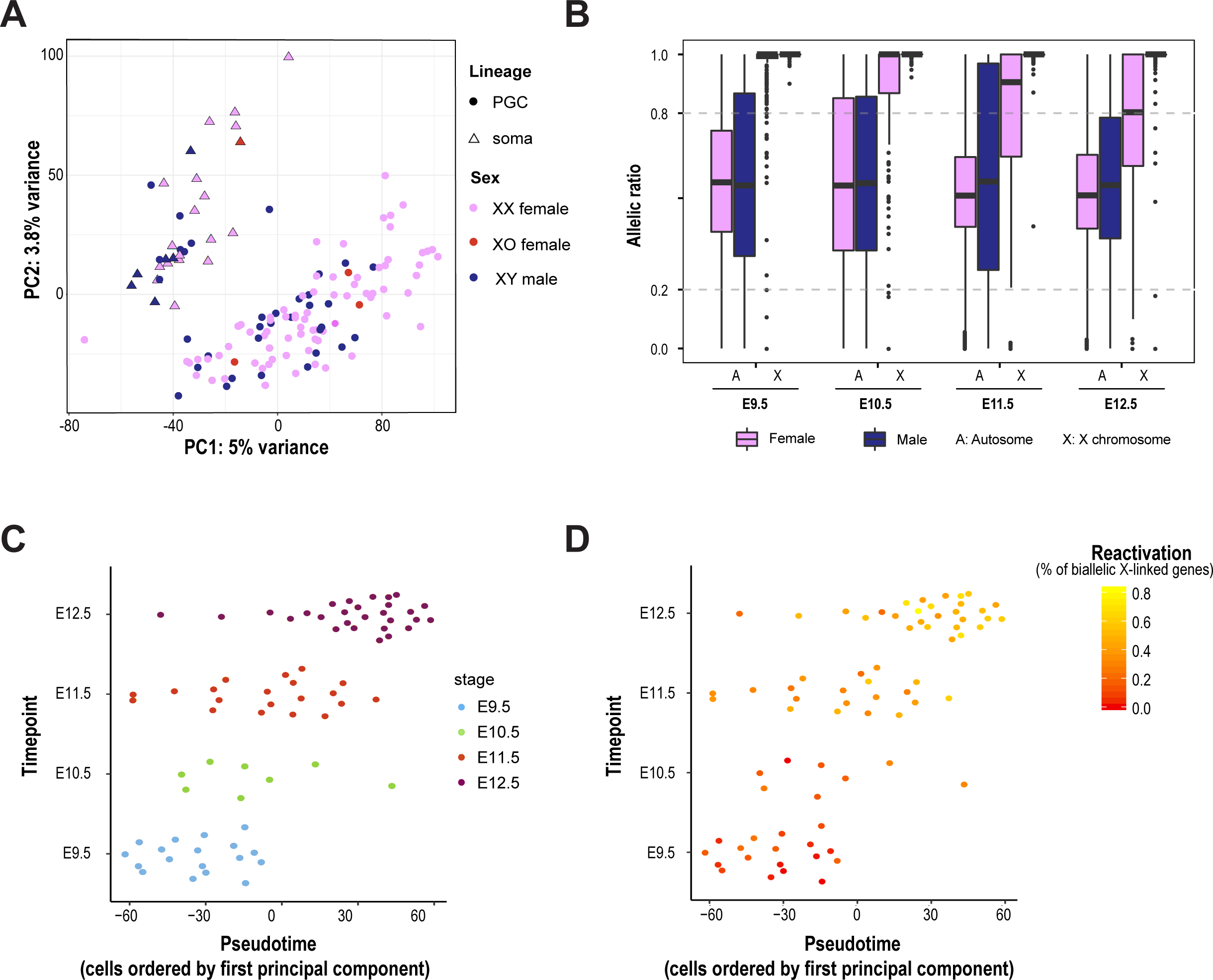
Progression of X-chromosome reactivation in developing PGC. (**A**) Principal component analysis (PCA) based on scRNA-seq data from PGC and soma between E8.5-E12.5. X-chromosome composition of each single cell is represented on the PCA, such as the XY male, XX female, and XO female cells. The XO cells originate from a single embryo and were found by serendipity. PCA is based on the 1 000 most variable genes, as described in Figure 1B. The details of each cell are listed in **Supplementary Table 1**. **(B)** Allele-specific expression ratios for genes on autosomes and X chromosomes in female and male single PGC from E9.5 to E12.5. The allelic ratio represents the number of reads mapped to Cast genome, divided by the total number of 129 and Cast reads. For X-linked genes, we measured the allelic ratio as the parental genome from which the Xi is originated, divided by the total number of 129 and Cast reads mapped for each gene (Xa counts / total Xa + Xi counts). A gene was considered biallelically expressed when 0.2< allelic ratio <0.8. Box plots represent medians (centre lines) with lower and upper quartiles (box limits). Whiskers represent 1.5× the interquartile range. Outliers are represented by dots. The number of cells analysed per stage, and the parental origin of the Xi, are shown in **Supplementary Table 1**. **(C)** and **(D)** Pseudotime representation of scRNA-seq data, based on the first principal component for female PGC between E9.5 and E12.5. In **(D)**, the percentage of reactivated X-linked genes per single cell is provided by a colour gradient as shown in the key. A gene is called reactivated if its allelic ratio is <0.8, which represents an expression from the Xi of at least 20 %

Furthermore, soma and PGC cells strongly clustered on PCA by lineage then developmental stage (1^st^ axis, **Figure 1B**). We then plotted male and female cells on a PCA for each developmental stage (**Extended Figure 2A-D**). Clustering by sex was confirmed from E12.5, based on gene expression (PCA, 1 000 most differentially expressed genes) in post-migratory E12.5 PGC, at the onset of sex gonadal differentiation (**Extended Figure 2D**). This was expected based on a previous large-scale scRNA-seq ^43^ and supported the quality of our database for further analysis.

### X-chromosome reactivation initiates progressively upon PGC development

To study X-chromosome reactivation, we analysed scRNA-seq in an allele-specific manner to determine the parental origin of the transcripts. SNPs from F1 hybrid embryos were used to map informative reads to either 129 or Cast genomes (see Methods). Each gene with informative SNP and expressed more than 2 Reads Per Retro-Transcribed length per million mapped reads (RPRT) were provided an allelic expression ratio (reads from Cast divided by total informative reads). The parental origin of Xi was then determined in each single cell based on the allelic ratio of all informative X-linked genes and the Xa allelic ratio calculated as reads mapped on active X (Xa) divided by total reads, in following analysis (**Supplemental Table 1,** see Methods). On the 94 female scRNA-seq, 64 cells carried an active X chromosome of Cast origin (68 %) and 30 cells of 129 origin (32 %). Despite random choice of Xi, we observed a strongly skewed silencing towards the 129 chromosome. This confirmed the X-chromosome controlling element (Xce) effect in F1 hybrid female mice. Indeed, it has been well documented that an F1 hybrid background could lead to skewed XCI depending on their Xce strength (Xce^a^<Xce^b^<Xce^c^<Xce^d^<Xce^e^ with the strongest Xce allele being the most resistant to silencing) ^49^. Using 129 and Cast strains associated with Xce^a^ and Xce^c^ respectively, we confirmed that the 129 chromosome was preferentially chosen to be silent in the expected proportions ^50^.

We then studied allelic expression of autosomal and X-linked genes during PGC development (**Figure 2B**). Autosomal genes were biallelically expressed (0.2< allelic ratio <0.8), with parity between 129 and Cast reads, in both males and females E9.5-E12.5 PGC. On the other hand, X-linked genes were strictly expressed from the Xa (allelic ratio > 0.8) in E9.5 female PGC, except for a few genes, presumably the escapees. Distribution of the X-linked gene allelic ratios was very similar in both female and male PGC, highlighting complete XCI in E9.5 PGC. From E10.5, X-chromosome reactivation initiated in female PGC (ratio<0.8) and progressed upon PGC development.

Because PGC could be heterogeneous in terms of developmental timing, even inside the same embryo, we decided to order the female PGC cells by pseudotime ordering (**Figure 2C**). We used the 1^st^ principal component of our PCA (**Figures 1B** and **2A**) to order female PGC by pseudotime ordering. Despite a few lagging cells, mainly at E11.5 and E12.5, most of the cells at the same developmental stage were clustered together, without a clear distinction of sex (**Figure 2C** and **Extended Figure 2E**). We then studied the percentage of reactivated X-linked genes in each female PGC (**Figure 2D**). We confirmed that Xi reactivation was progressive, from fully silent E9.5 PGC to highly reactivated E12.5 PGC, following PGC development (**Extended Figure 2E**).

Since the loss of *Xist* enrichment on Xi is the earliest known event during reactivation in the germline (based on IF/RNA-FISH) ^6, 10^, *Xist* expression levels were extracted in our scRNA-seq (**Extended Figure 2F**). We confirmed that *Xist* is not expressed in most PGC compared to female somatic cells, which restrains *Xist* expression and inactivates the X chromosome. In migratory E9.5 female PGC, X-chromosome reactivation has been initiated, despite the fact that most X-linked genes subject to XCI are still silenced.

Finally, to explore which pathways could drive Xi reactivation in the germline, we studied the correlation and anti-correlation between genome-wide gene expression and the percentage of reactivated genes (allelic ratio < 0.8) per female single cells (PGC and soma). Because X-chromosome reactivation occurs concomitantly to PGC development, we found that the best 2 correlated genes were germ cell-specific genes *Ddx4/Vasa* (R=0.78, q < 10^-12^) and *Dazl* (R=0.70, q < 10^-7^) ^1^. Gene ontology analysis of the top correlated genes (p < 0.001) revealed that chromatin modifiers involved in DNA methylation, gene silencing, and DNA modifications were overrepresented (**Extended Figure 3A**). This suggested that they may play a role in both PGC and Xi reprogramming.

### Different genes reactivate at different kinetics along the inactive X chromosome

Next, we investigated the kinetics of X-linked gene reactivation along the entire X chromosome. Heat maps of X-linked gene activity were generated for E9.5 to E12.5 female PGC (**Figure 3**). We focused on well-expressed genes to avoid confounding effects due to PCR bias and molecular loss in scRNA-seq method. Polymorphic genes expressed at RPRT>2 in at least 3 developmental stages were included in the heatmap and ordered by genomic position (**Figure 3 left**). E9.5 PGC displayed 92 % (164 out of 179) of monoallelically expressed genes (allelic ratio ≥ 0.8) and 8 % of biallelically expressed genes (0.2 < allelic ratio < 0.8), 8 of which are well-known escapees, such as *Kdm6a*, *Kdm5c*, *Ddx3x*, *Eif2s3x, Utp14a, Zrsr2, 1810030O07Rik, Pqbp1* ^51^. Interestingly, the proportion of biallelically expressed genes increased through the development of female PGC to reach 59 % (112 out of 189) at E12.5 PGC. This indicated the strong reactivation of X-linked genes during PGC development. However, an important portion of the X-linked genes was silenced at E12.5.

**Figure 3:**
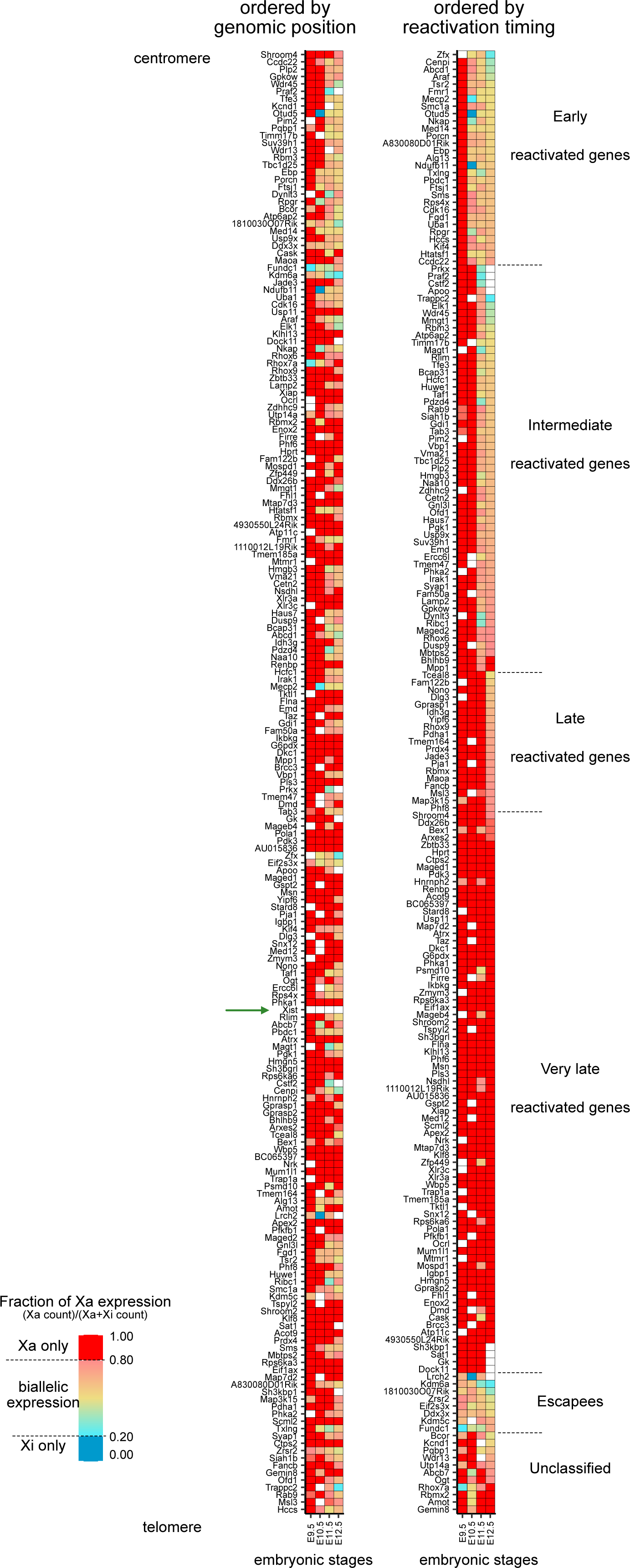
Kinetics of reactivation of X-linked genes over the entire X chromosome in developing PGC during reversal of random XCI. The mean of the allele-specific expression ratios, per embryonic stage, for each informative and expressed X-linked gene in female PGC are represented as heatmaps from E9.5 to E12.5, with strict monoallelic Xa expression (ratio >0.8) in red and strict monoallelic Xi expression (ratio <0.2) in blue. Color gradients is used in between these two values, as indicated in the key. Genes are ordered by genomic position (left) and reactivation kinetics class (right). *Xist* expression was always below RPRT < 2 and its genomic location has been added to the heatmap for information (green arrow). n = 198 informative X-linked genes, with a RPRT expression >2, expressed in at least 3 out of 4 developmental stages. White box, data not available (below threshold).

We classified the genes into different classes with respect to the timing of reactivation (**Figure 4A** and **Method, Figure 3 right**). At E9.5, all genes were silenced, except for the escapee class (n=8 out of 198 genes). Early genes (n=29 out of 198) were reactivated from E10.5, intermediate genes (n=55 out of 198 genes) from E11.5, and late genes (19 out of 198) from E12.5. At E12.5, 76 out of 198 genes were still silenced (monoallelically expressed from the Xa) and belonged to the very late-reactivated class.

**Figure 4:**
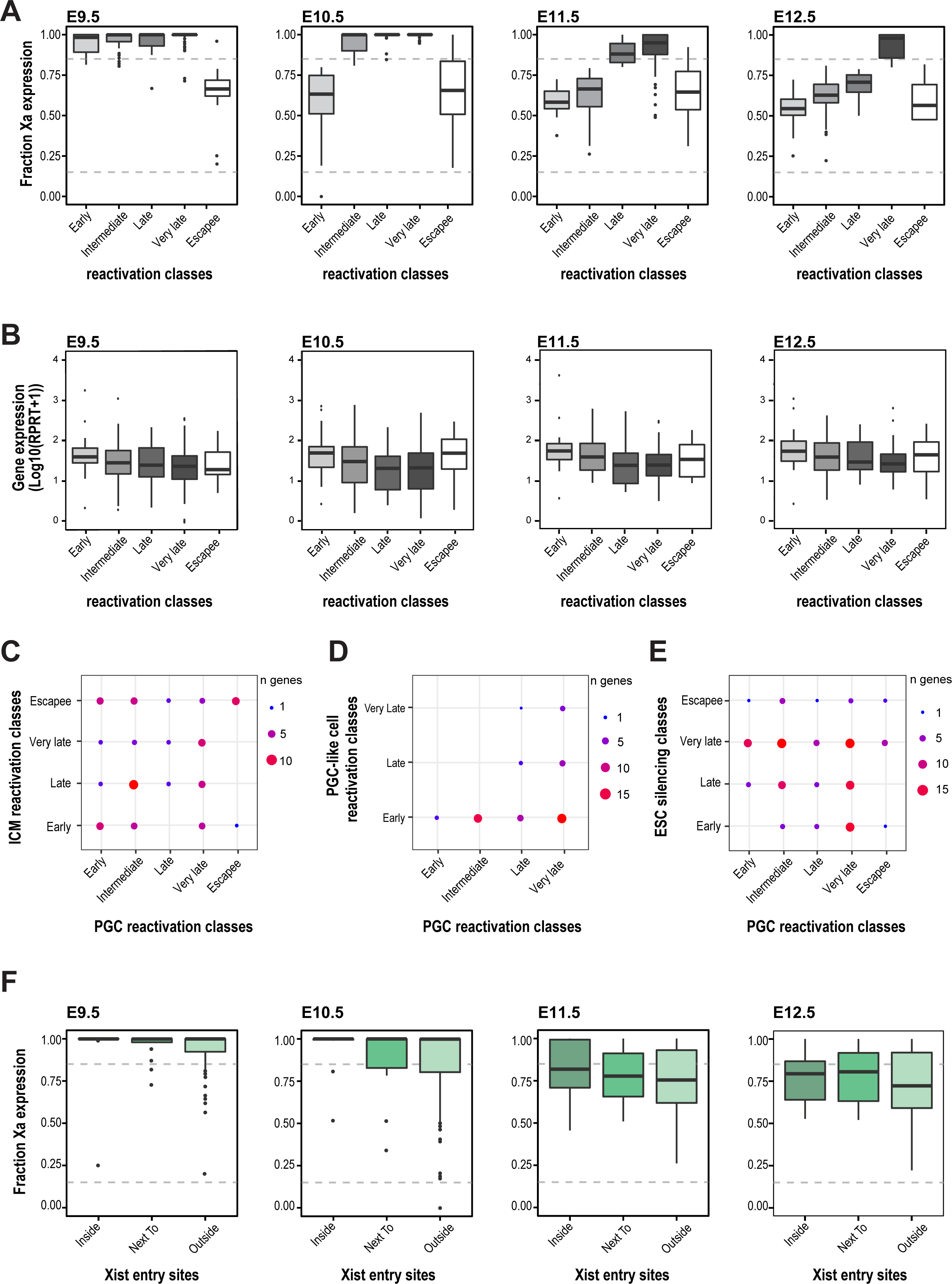
Differential timing of X-linked gene reactivation is associated with timing of silencing and chromosomal location in regards to *Xist* early sites. (**A**) X-linked genes are clustered based on their reactivation kinetics as early (expressed from the Xi at E10.5; allelic ratio <0.8 at E10.5), intermediate (expressed from the Xi from E11.5), late (expressed from the Xi from E12.5), very late (not reactivated at E12.5), and escapee (not undergoing XCI). The allelic ratio of each gene represents the fraction of Xa expression, with the number of reads mapped on the Xa genome divided by the total number of reads mapped. This is shown for stages E9.5 to E12.5 for all female PGC. n = 207 X-linked genes. Box plots are as in Figure 2B. Further information is provided in **Methods**. **(B)** Expression level of X-linked genes in the different reactivation-timing classes in female PGC (mean of each single gene). Expression of each gene represents the total number of reads mapped, normalized by the covered gene length and is represented at E9.5, E10.5, E11.5 and E12.5, as a function of the reactivation classes. Further information is provided in **Methods**. n = 207 X-linked genes. Box plots are as in Figure 2B. Comparison of reactivation classes between the inner cell mass of the blastocyst^29^ (**C**) and the *in vitro* PGC-like cell system^8^ (**D**). **€** Silencing classes of informative X-linked genes in mouse embryonic stem cells (mESC) compared with their reactivation classes in PGC^26^. **(F)** *Xist* ‘entry’ sites are regions of the X chromosome showing early accumulation of *Xist* RNA upon initiation of X-chromosome inactivation, and thought to be the closest to *Xist* locus in 3D spatial proximity. Allelic expression of X-linked genes classified on the basis of their relative position to *Xist* entry sites (as identified during XCI induction in ESC^18^): inside (TSS located inside a *Xist* entry site), next to (TSS located less than 100 kb away from an entry site) and outside (over 100 kb from an entry site). p=0.05 for E11.5 by Kruskall-Wallis test followed by Dunn’s correction. n = 207 X-linked genes. Box plots are as in Figure 2B. The numbers of cells analysed per stage is shown in **Supplementary Table 1**.

Differences in reactivation kinetics were not explained by different expression levels (**Figure 4B**). Early and escapee genes tended to be more highly expressed in E10.5 and E11.5, compared to still-silent genes. This could be explained by the fact that the Xi allele was also transcribed for early reactivated and escapee genes compared to the other classes of genes. Consistently, at E12.5, very late reactivated genes tended to be less expressed than those in the other classes.

A closer examination of the reactivation heatmap, ordered by genomic position, showed several regions of reactivation along the entire X chromosome. Because close genomic proximity to escapees could favour early reactivation in mouse iPSC ^31^, we tested the distance of our different reactivation class genes to the closest escapee (**Extended Figure 3B**). No link was detected between the differential reactivation kinetics and the distance from escapees in our *in vivo* female PGC. We then tested whether distance to *Xist* locus could be a significant parameter for reactivation kinetics (**Extended Figure 3C**). We found a strong bias for very late-reactivated genes to be localised closer to the *Xist* genomic locus compared to intermediate (p=0.02, KW test) and early (p=0.004, KW test) reactivated genes. Escapee were found further to the *Xist* locus (p=0.0045, KW test). We concluded that there was a strong correlation between close proximity to *Xist* locus and longer germline silencing.

### Context matters for the kinetics of X-linked gene reactivation

We aimed to test the consistency of the reactivation kinetics of X-linked genes in different developmental contexts (ICM vs. PGC). We compared the classes of reactivation that belong to common X-linked genes during imprinted Xi reactivation in ICM and random Xi reactivation in PGC (**Figure 4C**) ^29^. Few similarities were found, except for escapees in PGC, who also escaped imprinted XCI. We then compared the reactivation classes in PGC to a recent study on PGC-like cells (PGCLC) (**Figure 4D**) ^8^. The *in vitro* PGCLC model recapitulates early PGC specification, including Xi reactivation in the female cells. However, very few X-linked genes exhibited similar kinetics. *In vitro*, the cells underwent partial XCI; consequently, X-linked genes could be more prone to early reactivation. Late and very late reactivated genes in PGCLC could be genes that were properly silenced *in vitro* before undergoing reactivation. Thus, these genes had similar reactivation kinetics in our PGC *in vivo*. These data led us to hypothesize that late and very late reactivated genes in PGCLC, similar to PGC, could have been the first genes to be inactivated.

### Late-reactivated genes lay into *Xist* entry sites

To test the correlation between early XCI and late X reactivation, we compared the kinetics of reactivation in PGC to silencing in differentiating mES cells (**Figure 4E**) ^26^. We found that the early silenced genes belonged to the very late reactivation class in PGC. Being an early silent gene can influence the speed of reactivation a few days later.

Because the genes that resisted reactivation at E12.5, are closer to *Xist* locus (**Extended Figure 3C**) and could be the first silenced genes during random XCI (**Figure 4E**), this prompted us to question the relationship between X-linked gene reactivation and *Xist* entry sites. *Xist* entry sites are genomic regions of the chromosome that are the first bound by *Xist* RNA upon initiation of XCI^18, 20^. It is believed that *Xist* exploits the 3D conformation of the chromosome to first bind the regions in 3D spatial proximity to its transcription site and then initiate silencing before spreading across the entire chromosome ^20, 52^. Furthermore, we have previously shown that the genes lying inside these 3D accessible regions were more prone to early silencing *in vivo* ^16^. (**Figure 4F**). Here, we found that X-linked genes located within the *Xist* entry regions (Transcription Start Site TSS inside the predicted regions) showed more resistance to early reactivation than other genes. Genes outside the *Xist* entry sites showed the earliest reactivation and strongest allelic expression from Xi upon PGC development. Thus, the genes inside the *Xist* entry sites may correspond to genes that are more resistant to reactivation, carrying a stronger epigenetic memory of their silencing.*in vivo*.

### Resistance to reactivation could be partially explained by enrichment in chromatin repressive marks

Very late-reactivated genes appeared to correlate with the first silenced regions of the Xi. Because repeated sequences of the genome, mainly LINE-1 elements, have been proposed to help silence propagation and facultative heterochromatinization of the Xi, we tested the enrichment of transposons in and outside *Xist* entry sites (**Figure 5A**). We confirmed that the X chromosome is highly enriched in LINE-1 compared to control autosomal regions. However, we did not observe any significant enrichment of repeats inside *Xist* entry sites compared with the rest of the X chromosome, except for a slight enrichment of short interspersed nuclear elements (SINEs).

**Figure 5:**
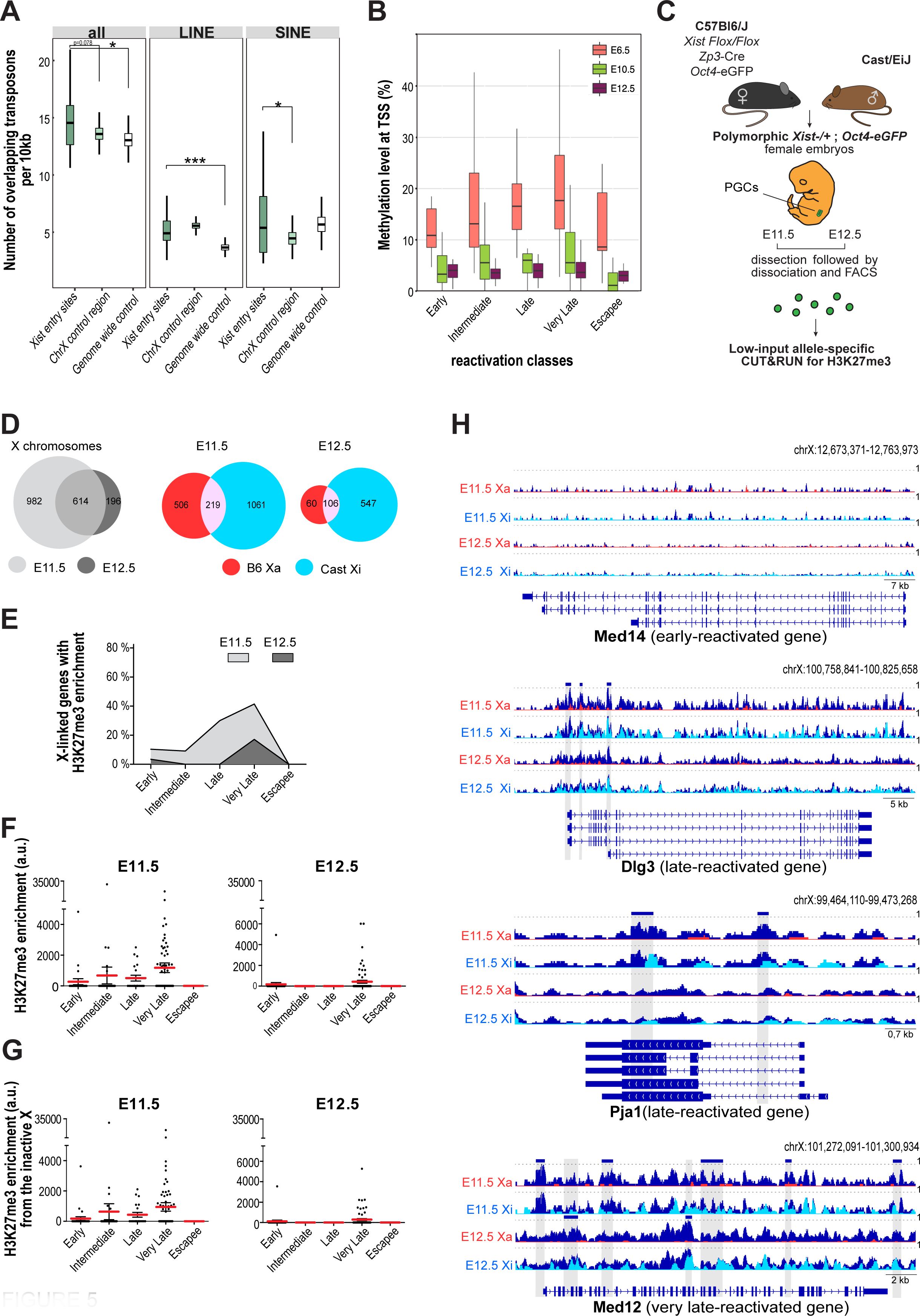
Contribution of repressive chromatin marks to resistance to early reactivation. (**A**) Number of total transposons, LINEs, and SINES, overlapping with *Xis*t entry sites, control regions of the X chromosome and control regions of autosomes. Set of control regions have been generated randomly 1000 times (see section **Methods**). A t-test was performed to compare number of repeats in *Xist* entry sites compared to controls.ass. **(B)** Whole-Genome Bisulfte of E6.5 female epiblast cells compared to public datasets of female PGC at E10.5 (DRA000607 in DDBJ database)^53^ and E12.5 (GSE76971 in GEO database)^54^. DNA methylation level were estimated using a window of -1kb to +100bp to the TSS for each gene. **(C)** Schematic illustration of the mouse breeding between C57Bl6/J *Xist flox/flox*; *Zp3*-Cre; *Oct4-*eGFP females and Castaneus males in order to obtain female polymorphic embryos, with non-random XCI and fluorescent PGC. PGC were isolated by FACS with the Oct4-eGFP reporter at E11.5 and E12.5. Following PGC sorting, low-input CUT&RUN was completed for H3K27me3 marks (see **Methods**). **(D)** Distribution of H3K27me3 CUT&RUN peaks in E11.5 and E12.5 female PGC. The Venn diagram shows H3K27me3 broad domain overlapping in X chromosomes between E11.5 and E12.5 female PGC (left), and between the active (B6, Xa) and inactive (Cast, Xi) chromosomes, right. **(E)** Percentage of early (n=29), intermediate (n=55), late (n=20), very late (n=70), and escapee (n=8) X-linked genes significantly enriched in H3K27me3 repressive marks in both E11.5 and E12.5 female PGC (at least one H3K27me3 broad domain per gene). **(F)** H3K27me3 enrichment (fold change compared to IgG and normalized by library size and peak length) in the different X-linked gene reactivation classes. Kruskall Wallis test followed by Dunn’s post hoc test was performed to compare all classes. Each point represents a gene, in red the mean +/-sem. E11.5: p-value < 0,0001, early and intermediate versus very late and E12.5: p-value = 0,0020 intermediate versus very late. **(G)** enrichment (fold change compared to IgG and normalized by library size and peak length) weighted by the d-score (allelic ratio, see **Methods**) at E11.5 (p-value = 0,0004, early and intermediate versus very late), E12.5 (p-value = 0,0038, intermediate versus very late). Statistical test (mean, SEM). Each point represents a gene, in red the mean +/-sem. E11.5: p-value < 0,0001, early and intermediate versus very late and E12.5: p-value = 0,0020 intermediate versus very late. **(H)**. Integrative Genomics Viewer plot of representative genes from early, late and very late reactivation classes. Tracks depict global and allele-specific H3K27me3 enrichment for *Med14* early reactivated gene, *Dlg3* and *Pja1* late reactivated genes and *Med12* very late reactivated gene. Global enrichment tracks are in dark blue and allele-specific tracks are overlaid with global enrichment tracks (Cast Xi reads in light blue; B6 Xa reads in red). Dark blue boxes and highlighted grey area are significant H3K27me3 broad domains. Location is given in mm10, with gene isoforms extracted from Integrative Genome viewer and UCSC.

Different chromatin environments could explain the differential kinetics of reactivation of X-linked genes in the PGC, as previously demonstrated in the ICM of blastocysts ^29^. Both repressive histone marks, such as H3K27me3 and DNA methylation, are enriched on the Xi upon random XCI ^14^. We studied DNA methylation by whole-genome bisulfite sequencing (WGBS) in female E6.5 epiblasts, when XCI takes place, and in publicly available datasets of female E10.5 ^53^ and E12.5 ^54^ PGC (**Figure 5B**). However, because XCI is random, the population of PGC was heterogeneous in terms of parental origin of the Xi (*i.e.* 50 % of the cells silence the paternal X chromosome and 50 % the maternal chromosome), and cell population*-*based assays on mosaic female embryos are not informative for deciphering between Xa and Xi. Thus, we considered for the following analysis that a significant enrichment of DNA methylation at X-linked gene promoters had a higher probability of coming from the Xi rather than the Xa. Very late reactivated genes showed slight enrichment in DNA methylation at their TSS compared to early-reactivated genes at E6.5. From E10.5, the DNA methylation erasure was nearly complete, and differences seemed to be lost (**Figure 5B**). Although very late reactivated genes were initially more enriched in DNA methylation than the early reactivated ones, it is not clear why these genes still resist reactivation at E12.5, when DNA demethylation is complete.

We then decided to study the repressive histone mark H3K27me3, which is enriched on the inactive X chromosome and confers resistance to early reactivation of some X-linked genes in the ICM ^29^. To overcome the mosaicism in the PGC population and decipher between Xa and Xi, we used triple transgenic female mice (*Xist ^flox/flox^: Zp3-Cre; Oct4(ΔPE)eGFP*, on a C57Bl6/J genetic background) crossed with Cast males (**Figure 5C**). This allowed us to collect polymorphic female embryos and sort pure PGC populations based on GFP (under the promoter control of the pluripotency factor OCT4) with non-random X inactivation. The *Xist* deletion occurred in the maternal germline and allowed the transmission of a Xist^KO^ B6 allele, which cannot be inactivated; Xi being always the Cast allele. Low-input allele-specific CUT&RUN against H3K27me3 marks was performed on sorted GFP+ female PGC at E11.5 and E12.5 (**Figure 5C**). Statistically enriched broad domains of H3K27me3 were identified on X chromosomes (**Figure 5D**). We found a higher number of peaks at E11.5 on the X chromosome than at E12.5, in which most of the peaks were conserved from E11.5. The reads were then mapped to the parental genomes to determine the parental origin of the reads. As expected, most peaks were found on the inactive X chromosome at both E11.5 and E12.5 (**Figure 5D** and **Extended Figure 4).** We then intersected H3K27m3 enrichment with reactivation classes. Enrichment in K27me3 was mainly found at E11.5, (20 % of the late-reactivated and 45 % of very late-reactivated genes). At E12.5, H3K27me3 enrichment was lost in late genes when they were transcriptionally reactivated (**Figure 5E).** The enrichment level is also more important in very late reactivated genes, with most of the signal coming from the inactive Cast chromosome (**Figures 5F-G).** In contrast, early-reactivated genes were depleted in H3K27me3, in accordance with their biallelic expression status. Tracks showing global and allele-specific H3K27me3 enrichment were produced for the early reactivated gene *Med14*, late *Dlg3* and *Pja1* genes, and very late *Med12* (**Figure 5H**). H3K27me3 enrichment was not detected at *Med14* gene location. Both *Dlg3* and *Pja1* were statistically enriched at E11.5, but not at E12.5, once they were reactivated. Finally, *Med12* gene was strongly enriched at both E11.5 and E12.5, and showed no reactivation at E12.5, based on scRNA-seq. We observed that some late reactivated genes could carry an epigenetic memory of their silencing (H3K27me3), which was lost concomitantly to transcriptional reactivation.

## Discussion

In this study, we performed a comprehensive allele-specific single-cell transcriptomic analysis of migratory and colonizing female PGC, combined with low-input epigenomics. We demonstrated that female early PGC carry a fully inactive X chromosome, which undergoes progressive reactivation in parallel with PGC development. Although it was previously established that the inactive X chromosome experiences transcriptional reactivation in PGC, our study provides the first detailed map of X-chromosome activities *in vivo*. We showed that different genes followed different kinetics of reactivation along the chromosome, with early versus late reactivated genes. We provide evidence for the involvement of genomic location, 3D spatial proximity to the *Xist* locus, and H3K27me3 chromatin modification in resistance to early reactivation. Together, these investigations open a way for a better understanding of the *in vivo* requirements for female epigenetic reprogramming in general, stem cell biology, and reproduction.

X-chromosome reactivation occurs during the reprogramming of female primordial germ cells. Consequently, this leads to an excess of X-linked gene products in female PGCs compared to males ^10^. In mice, this happens transiently at the onset of sex-specific gonadal differentiation (E9.5-<E15.5) in the female germline and could be crucial for normal gonadal development and meiosis, as well as for sex-specific reprogramming. This could promote sexual dimorphism. Patients with sex-chromosome aneuploidy, such as Turner 45,XO, and Klinefelter 47, XXY syndromes, often present infertility and hypogonadism. Maternally inherited sex-chromosome aneuploidy could arise from the presence of a non-reactivated X, potentially detrimental to homologous chromosome pairing and segregation in meiosis ^12^. Our transcriptomic analysis of XX female, XO female, and XY male PGC highlighted differentially expressed genes, which could be involved in germline formation and/or sex-specific differences. The X chromosome is enriched in factors required for oogenesis (*e.g.* Fmr1, Zfx) and chromatin modifications and transcription (e.g., histone demethylases Kdm6a, Kdm5a, mediator complex Med14). Importantly, we showed in this study that all these factors are either not subject to XCI or are early-reactivated in female PGC. *Med14* codes for a co-unit of Mediator complex, involved in transcription regulation, and is prone to early reactivation in both mouse iPS ^31^ and human breast cells depleted for XIST ^55^. *Smc1a* gene has recently been shown to be involved in the remodelling and reactivation of Xi in mouse iPSC ^56^. *Kif4* knock-down in oocytes is detrimental to meiosis ^57^. *Zfx* is a well-known dose sensitive gene. Its absence in mouse leads to infertility owing to the reduced number de germ cells ^58^. Together, this raises the importance of studying these genes with dosage imbalance in PGC. An appropriate dosage of X-linked genes, whose functions are linked to chromatin processes, transcription, and gametogenesis, could be important for female gametogenesis and X-chromosome reactivation.

Our detailed mapping of X-linked gene reactivation kinetics highlights differential behaviours along the entire X chromosome. The differential kinetics of reactivation are dependent on the developmental context ^11, 29, 31^. Understanding the resistance to early reactivation and the underlying mechanism is important for understanding epigenetic reprogramming on the X-and genome-wide levels. Surprisingly, we showed that 40 % of the Xi remained silent at E12.5. These genes lay into regions in close 3D proximity to *Xist* locus (*Xist* entry sites) and could be the first genes to be silenced upon XCI. We believe that these first-silenced genes could be the first targets of *Xist* because of their 3D accessibly^52^. They would become less accessible to the transcriptional machinery and are more enriched in repressive marks, carrying an epigenetic memory of their silencing. In support of this hypothesis, our results indicated that the latest reactivated genes at E12.5 are still enriched in H3K27me3 on their silent allele. These repressive marks are lost concomitantly with gene reactivation.

In conclusion, in PGC, we observed a reactivation dependency on *Xist* RNA loss, DNA methylation level, enrichment in transposable elements, and the proximity of X-linked genes to the first regions coated by *Xist* RNA. Together, these investigations open a way for a better understanding of the in vivo requirements for female epigenetic reprogramming in general, stem cell biology, and reproduction.

## Methods

### Mouse husbandry

The care and use of animals are strictly applying European and National Regulation for the Protection of Vertebrate Animals used for Experimental and other Scientific Purposes (Directive 2010/63/EU and French decree R.214-103). All husbandry and experiments involving mouse scRNA-seq were authorised by the UK Home Office Project Licenses PPL80/2637 and PE596D1FE and were carried out in a Home Office designated facility (Welcome Trust Cancer Research Gurdon Institute). Chromatin experiments were authorized by the French ethics committee number 36 under agreement F3417216 and carried out in the pathogen-free Animal Care Facility of IGMM (facility licence #G34-172-16). Researchers carrying out regulated procedures on living mice held a personal licence from either the UK (C.L., M.B., and M.A.S.) or France (C.R., M.B., K.C., and D.B.).

Mice were housed under a 12h light/12h dark cycle at 22 ± 2 °C ambient temperature, with free access to food and water. All embryos were derived from natural mating. Noon on the day of observation of the vaginal plugs was scored as embryonic day (E) 0.5. Embryos were harvested every 24 h between E8.5 and E12.5. Collected embryos were included in the analyses only if they showed normal morphology according to their developmental stages. No statistical method was used to determine the sample size.

Male and female hybrid embryos were obtained by breeding *Mus musculus domesticus* 129S1/SvImJ *Stella-eGFP* transgenic line^39^ (at E8.5) or *Mus musculus domesticus* 129S1/SvImJ *GOF-△PE-18* transgenic line^40^ (from E9.5 onwards) with *Mus musculus castaneus* (CAST) (**Figure 1A**). *Xist* ^-/+^ *Zp3-*CRE; *Oct4-*eGFP female embryos (**Figure 5C**) were obtained by mating *Mus musculus domesticus* C57Bl6/J *Xist*^flox/flox^; *Zp3-* CRE; *Oct4-*eGFP females^17, 59, 60^ with *Mus Musculus Castaneus* males.

### Sexing of the embryos

The sex of the embryos was characterized based on the morphology of the gonads from E11.5. Before E11.5, sex was characterized in single-cell RNA-seq datasets by studying the expression of *Xist* and Y-linked genes, as well as the presence or absence of polymorphisms (SNPs) on the X chromosome. For the WGBS, and CUT&RUN experiments, the sex of the embryos was determined by PCR using genomic DNA and Ube1 primers (Ube1-Forward *TGGATGGTGTGGCCAATG*; Ube1-Reverse *CACCTGCACGTTGCCCTT*).

### Collection of PGC

After dissection of embryos at the location of the PGC, according to embryonic stage, and sexing of the embryos, samples were resuspended in 200 µL of 0.25 % trypsin and incubated at 37 °C for 3 min. Trypsin was inactivated with serum and a single-cell solution was obtained by vigorous up-and-down. For scRNA-seq experiments from E10.5, cells were manually picked based on their GFP and size and washed in PBS-acetylated BSA (**Figure 1A**). For the other stages and CUT&RUN experiments, cells were collected by fluorescence-associated cell sorting (FACS ARIA© and S3e Cell Sorter Bio-Rad©) and processed quickly for scRNA-seq or low-input CUT&RUN.

### Single cell RNA sequencing and bioinformatic analysis

Single PGC were washed thrice with PBS/acetylated BSA (Sigma) before being manually transferred within the minimum amount of liquid into PCR tubes. We either directly prepared the cDNA amplifications or kept the single cells at − 80 °C (less than 2 months) for future preparation. Poly(A)^+^ mRNA extracted from each single cell was reverse-transcribed from 3’-UTRs and amplified according to a previously described protocol^42, 61^. Care was taken to process only embryos and PGC of the highest quality based on morphology and amplification yield. A total of 140 single cells were processed and quality control (QC) was performed as previously described in ^16^.

Single-cell libraries were prepared from 137 samples that passed QC, according to the manufacturer’s protocol (Illumina). Sequencing to produce single-end 50-bp reads was then performed on an Illumina HiSeq 4000 instrument (**Supplementary Table 1**).

Quality controls, filtering of raw data, mapping, and SNP calling have been described previously ^8, 9^. Briefly, the mouse mm10 genome was downloaded from Sanger database. To study allele-specific gene expression, reads were processed according to Borensztein et al^16^. SNPs between the 129 and Cast strains were extracted from the VCF file and used to reconstruct the Cast genome. After the removal of the common exonic SNPs between *Xist* and *Tsix*, 20,220,776 SNPs were retained. The number of paternal and maternal reads were counted at each SNP position. The threshold used to call a gene informative was five reads mapped per single SNP, with a minimum of eight reads mapped on SNPs per gene, to minimize disparity with low-polymorphic genes. The allele-specific origin of the transcripts (allelic ratio) was calculated as the total number of reads mapped to the Cast genome divided by the total number of reads for each gene: allelic ratio = Cast reads/(Cast + 129) reads. For X-linked gene, we modified the allelic ratio to: allelic ratio = Xa reads/(Xa + Xi) reads. In the case of a 129 Xi, the allelic ratio became 1-[Cast reads/(Cast+129) reads]. Genes were thus classified into two categories: inactivated genes: allelic-ratio value ≤0.20 or ≥0.80, and biallelically expressed genes: allelic-ratio value >0.20 or <0.80.

*Estimation of gene expression levels.* Given that our RNA reverse transcription allowed sequencing only up to an average of 3 kb from the 3′ UTR, half of the expressed genes were only partially covered (less than 50% of the gene size on average). To estimate transcript abundance, read counts were normalized based on the amplification size of each transcript (RPRT) rather than the size of each gene (RPKM) (see details in Borensztein et al.^42^).

*Principal component analysis, hierarchical clustering, and differentially expressed genes in volcano plots.* Only genes with an RPRT value >1 in at least 25% of the single cells of at least one developmental stage (with a minimum of two cells) were retained for downstream analysis, as previously described in ^16^. With the Benjamini–Hochberg correction, genes with an adjusted *P* value lower than α = 0.05 were called as differentially expressed.

*Heatmap generation for X-chromosome allelic gene expression.* For allelic ratio heatmaps, data from informative genes were analysed at each developmental stage only if the gene was expressed (RPRT >2) in at least 25% of the single blastomeres (with a minimum of two cells) (**Figure 3**). To follow the kinetics of expression, we focused only on genes expressed in at least three different stages. The mean allelic ratio of each gene is represented for the different stages of the female PGC.

*Global gene expression correlation with X-chromosome reactivation.* Correlation and anti-correlation between gene expression levels (autosomes and X chromosomes) and the percentage of X-linked gene reactivation (allelic ratio <0.8 for X-linked genes) were measured using Pearson’s correlation and Benjamini–Hochberg correction. Gene ontology analysis was performed for the top-correlated genes (q-value <0.05). Genes with RPRT < 2 were considered to be unexpressed (RPRT = 0 and allelic ratio = NA).

*Definition of X-linked gene reactivation classes.* We automatically assigned X-linked genes to the reactivation classes.

– Early reactivation: expressed on both chromosomes at stage E10.5. Allelic ratio >= 0.8 or NA at E9.5; allelic ratio < 0.8 at E10.5; allelic ratio < 0.8 at E11.5; allelic ratio < 0.8 or NA at E12.5.

– Intermediate reactivation: expressed on both chromosomes at stage E11.5. Allelic ratio >= 0.8 or NA at E9.5; allelic ratio >= 0.8 or NA at E10.5; allelic ratio < 0.8 at E11.5; allelic ratio < 0.8 or NA at E12.5.

– Late reactivation: expressed on both chromosomes at stage E12.5. Allelic ratio >= 0.8 or NA at E10.5; allelic ratio >= 0.8 at E11.5; allelic ratio < 0.8 at E12.5.

– Very Late reactivation: silenced at stage E12.5. Allelic ratio >= 0.8 at E10.5 Allelic ratio >= 0.8 at E11.5 and E12.5 OR allelic ratio < 0.8 at E11.5 and >= 0.8 at E12.5.

– Escapees: always biallelic. Allelic ratio < 0.8 at all stages (**Figure 4A**).

### Low-input CUT&RUN

After PGC were collected, the cells were directly pelleted at 4 °C for 5 min. The CUT&RUN protocol was modified from Skene *et al*^64, 65^ and Dura *et al*^66^ to accommodate a low number of PGC (5 000-15 000 per sample, **Supplementary Table 1**). Briefly, cells were split according to the number of required antibody profiles, and Nuclear Extraction Buffer (20 mM HEPES-KOH, 10 mM KCl, 0.5 mM spermidine, 0.1 % Triton X-100, 20 % Glycerol, Complete EDTA-free protease inhibitor cocktail) was gently added to the cell solution and incubated on ice for 5 min. Cells and concanavalin A beads were incubated for 10 min at room temperature (RT) on a rotating wheel. Cells were then collected on magnets, resuspended in Blocking Buffer (20 mM HEPES-KOH, 150 mM NaCl, 0.5 mM spermidine, 0.1 % BSA), 2 mM EDTA, and 1× Complete EDTA-free protease inhibitor cocktail), and incubated at RT for 5 min. Cells were then washed and incubated with H3K27me3 antibody (1:200 dilution, Cell signalling 36B11#9733, control with IgG rabbit Sigma) for 2h30 at 4 °C on a rotating wheel and then washed twice. Samples were then incubated with 1:400 Protein A-MNase fusion protein (gift from the Dominique Helmlinger lab, CRBM France) for 1 h at 4 °C followed by two washes. Cells were then resuspended in 150 µL Wash Buffer and cooled in an ice-water bath for 5 min before the addition of a final concentration of 100 mM CaCl_2_. Targeted digestion was performed for 30 min on ice. The samples were then incubated for 20 min at 37 °C to release the cleaved chromatin fragments. After centrifugation at 16,000 × *g* for 5 min, supernatants were transferred to new low-binding tubes. Following the addition of 20 % SDS and 20 mg ml^−1^ Proteinase K, the samples were incubated 30 min at 70 °C. DNA was purified using phenol/chloroform, followed by chloroform extraction and precipitation with 20 mg ml^−1^ glycogen and three volumes of 100 % ethanol at 20 °C. The DNA pellet was washed with 85% ethanol, centrifuged, and resuspended in low Tris-EDTA.

Library preparation was performed according to manufacturer’s instructions (NEBNext^®^ Ultra™ II DNA Library Prep Kit for Illumina) using the following modified library amplification program: 98 °C for 30 s (98 °C for 10 s, 65 °C for 15 s) × 15 cycles, 65 °C for 5 min), hold at 4 °C. Average library size and quality control were performed using a Fragment Analyzer (High Sensitivity NGS kit) and qPCR (Roche Light Cycler 480). CUT&RUN libraries were sequenced on a NovaSeq 6000 (Illumina) from Biocampus, MGX platform, using a paired-end 150-bp run. E11.5, and E12.5 H3K27me3 CUT&RUN were performed on one replicate of five and two pooled female embryos, respectively.

### Allele-specific CUT&RUN bioinformatic analysis

FastQC (v0.11.9) and MultiQC (v1.13)^67^ were used to control CUT&RUN data quality. UMIs were used to confirm the homogeneous yield of library amplification and sequencing for each sample. Paired-end reads with at least one undefined UMI were discarded using Cutadapt v4.1^68^ and seqkit v2.3^69^. The reads were then trimmed using Trim Galore(v0.6.6^16^) (https://www.bioinformatics.babraham.ac.uk/projects/trim_galore/70; options “--length 20 –illumina – 2colour 20”). Trimmed reads were mapped with Bowtie2 (v2.4.2)^71^ (options “--end-to-end –very-sensitive –reorder”) to the mm10 reference genome, modified as followed: autosomal, sexual and mitochondrial chromosomes are kept, and 20 668 274 SNPs positions (0.76% of genome size) related to *Mus musculus Castaneus* strain are N-masked using SNPsplit (v0.6)^72^ (https://ftp.ebi.ac.uk/pub/databases/mousegenomes/REL-1505-SNPs_Indels/strain_specific_vcfs/CAST_EiJ.mgp.v5.snps.dbSNP142.vcf). SAMtools (v1.11)^73^ was used to sort and convert the data formats. PCR duplicates were removed using GATK MarkDuplicates, and reads were unmapped with or without primary alignment discarded (SAMtools options “-F 0×04 -F 0×100 -F 0×800”). The coverage for each developmental stage was calculated from the bam file using the deepTools bamCoverage tool (v3.5.1, normalization by scale factor, bin=10 nt)^74^.

Peak calling of H3K27me3 histone marks was performed for E11.5 and E12.5 female samples using immunoglobulin G (IgG) as input for each stage (E11.5 IgG from male PGC, E12.5 IgG from female PGC) with MACS2 (v2.2.7.1, options “-f BAMPE –broad –broad-cutoff 0.1”)^75^.

Allele-specific reads (*i.e.* reads from B6: Xa and Cast: Xi) were sorted using SNPsplit. The d-score parameter (reads Cast / reads B6 + reads Cast]) was calculated using featureCounts from the Rsubread R package(v2.12.3)^76^. H3K27me3 enrichment heatmaps show the log2 fold change in the coverage difference between the H3K27me3 mark and the IgG control for each developmental stage, normalized by the sequencing depth using deepTools bamCompare (bin= 10 nt). computeMatrix and plotHeatmap.

### Whole Genome Bisulfite sequencing of female Epiblast

C57Bl6/J Epiblasts were manually dissected from extra-embryonic tissues of E6.5 embryos, followed by sex determination by PCR on the extra-embryonic tissue (see section sexing of the embryos). Whole-Genome Bisulfite sequencing libraries from 2 E6.5 female replicates were prepared as described by Smallwood et al^77^. The WGBS was analysed as described previously^78^. Briefly, reads generated in this study or recovered from the available datasets were treated as follows: the first eight base pairs of the reads were trimmed using the FASTX-Toolkit v0.0.13 (hannonlab.cshl.edu/fastx_toolkit/index.html). Adapter sequences were removed with Cutadapt v1.3 (code.google.com/p/cutadapt/)^68^ and reads shorter than 16 bp were discarded. The cleaned sequences were aligned to the mouse reference genome (mm10) using Bismark v0.12.5 70 with Bowtie2-2.1.0 71 and the default parameters. Only the reads that mapped uniquely to the genome were conserved. Sequencing statistics can be found in the Figure Legend and/or the main text. Methylation calls were extracted after duplicate removal. Only CG dinucleotides covered by a minimum of ten reads were conserved for the remainder of the analysis.

### DNA methylation and transposon data analysis

DNA methylation data at E10.5 and E12.5, were downloaded from DRA000607^53^ and GSE76971^79^ respectively. The raw data were cleaned using Trim Galore v0.4.4^70^. The cleaned reads were aligned to the mouse reference genome assembly (GRCm38/mm10) using Bismark v0.18.2^80^ with Bowtie2-2.2.9^71^ allowing for one mismatch in the seed alignment. Only reads that mapped uniquely to the genome were retained, and methylation calls were extracted after duplicate removal, considering only CpG dinucleotides covered by a minimum of five reads.

For the control regions of transposon enrichment, a set of 25 random regions (number of Xist entry sites), with the same length as the median *Xist* entry sites) was bootstrapped 1 000 times either genome-wide or on the X chromosome, using the R package regioneR (v1.10.0, Gel B, Diez-Villanueva A, Serra E, Buschbeck M, Peinado MA, Malinverni R (2016). “regioneR: an R/Bioconductor package for the association analysis of genomic regions based on permutation tests.” *Bioinformatics*, **32**(2), 289-291). The number of retrotransposons overlapping *Xist* entry sites and random regions was calculated and normalized to a 10 kb window. DNA methylation levels were estimated using a window of -1 kb to +100 bp from the Transcriptional Start Site (TSS) for each gene. In all the cases, a permutation test was performed using the RegioneR package.

Statistics.

Statistical significance was evaluated using Kruskal–Wallis followed by Dunn’s correction and t-tests. *P* values are provided in the figure legends and/or the main text.

## Data availability

All sequencing data will be deposited in GEO and made publicly available after publication of this article in a peer-reviewed scientific journal.

## Authors’ contribution

M.B. conceived the study with M.A.S. and performed scRNA-seq experiments. C.R. performed chromatin analysis, handled mouse colonies, and collected embryos with the help and supervision of K.C.. L.S., E.B., and D.Z. performed bioinformatics analysis, supervised by N.S. and M.B.. A.T. performed the repeats and WGBS bioinformatics analysis. C.L. contributed to the animal husbandry and sample collection. M.W. and D.B. conducted WGBS experiments. M.B., M.A.S., and D.B. secured the funding. M.B. wrote the original manuscript. with inputs from co-authors.

## Acknowledgments

We acknowledge the Surani, Bourc’his, and Heard labs for insightful discussions, Kay Harnish, Cambridge Stem Cell Institute Genomics Facility, and the MGX-Montpellier GenomiX platform for advice and deep sequencing of the libraries. We thank the pathogen-free barrier animal facility of Gurdon Institute and the PCEA and ZEFI of IGMM, UMR5535, and Montpellier. We are grateful to Joan Barau, Dominique Helmlinger, and Lorraine Bonneville for their advice and reagents regarding the low-input CUT&RUN.

This work was supported by a CNRS-INSERM ATIP-Avenir grant, the FRM AJE202005011598 and an ANR (the French National Research Agency) under the "Investissements d’avenir" programme with the reference ANR-16-IDEX-0006 » to M.B.; Wellcome Trust funding 096738 and 092096 and Cancer Research UK program C6946/A14492 to M.A.S., by a PhD fellowship from La Ligue nationale contre le cancer to C.R. and FRM SPE20150331826 and a Marie Sklodowska-Curie Individual Fellowship (H2020-MSCA-IF-2015—no. 706144) to M.B. MGX acknowledges financial support from France Génomique National infrastructure, funded as part of “Investissement d’Avenir” program managed by Agence Nationale pour la Recherche (contract ANR-10-INBS-09).

---

**Extended Figure 1.**
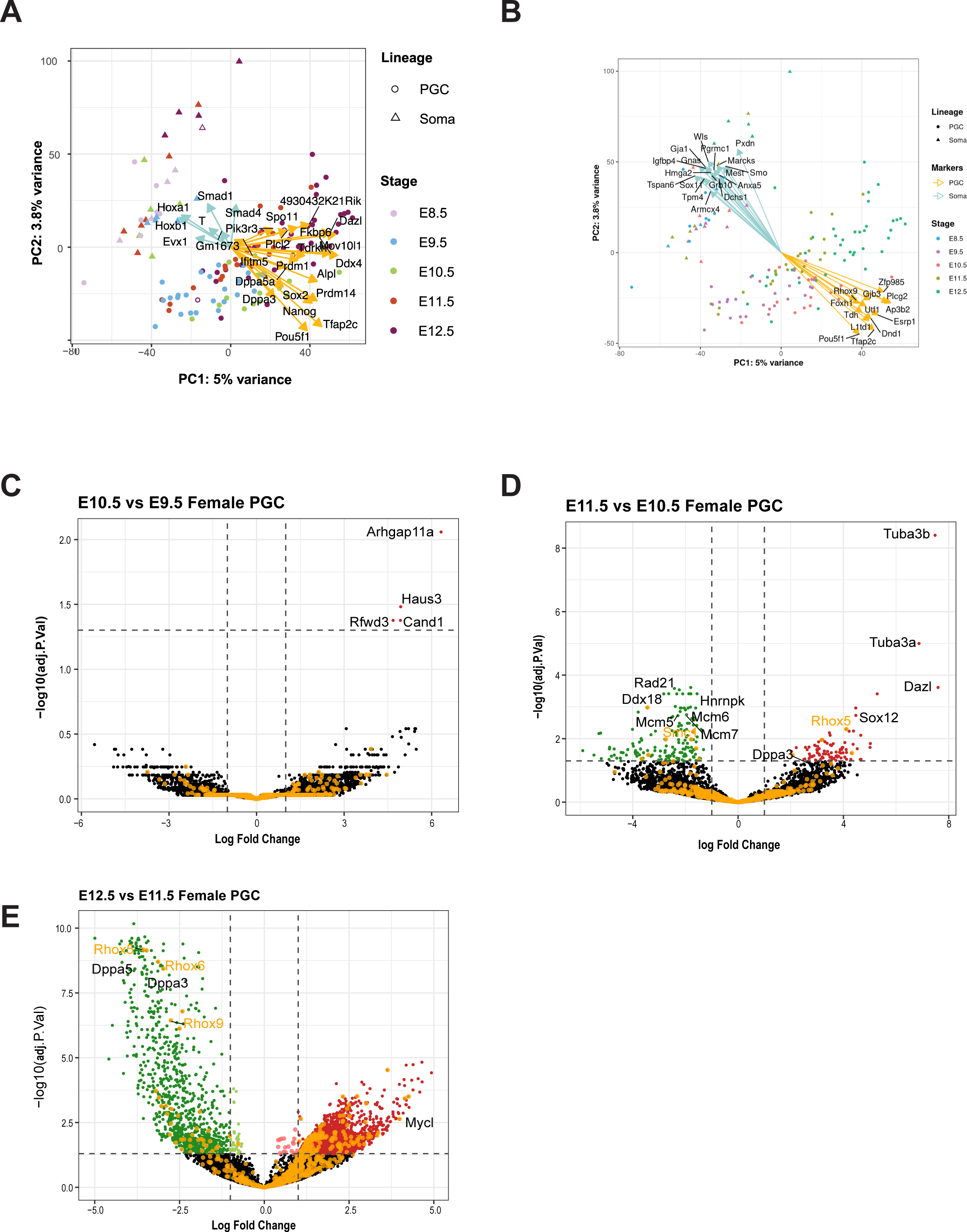
Differential gene expression upon female PGC development and key markers of soma and PGC. **(A)**. Principal Component Analysis (PCA) based on 26 known markers of PGC or soma (Figure 1C and D). **(B)** PCA of the 30 most differentially expressed genes (DEGs) that contributed to lineage segregation. **(C-D)** Volcano plots represent differentially expressed genes (DEG) between the two developmental stages of female PGC. A few transcriptional changes have been observed in migratory PGC. Changes arise once PGC colonize the gonads. Some examples of DEG are highlighted. Red dots represent upregulated genes, and green dots represent downregulated genes. X-linked genes are shown in orange. They showed a statistically greater enrichment in upregulated genes at E12.5, compared to E11.5, owing to X reactivation.

**Extended Figure 2.**
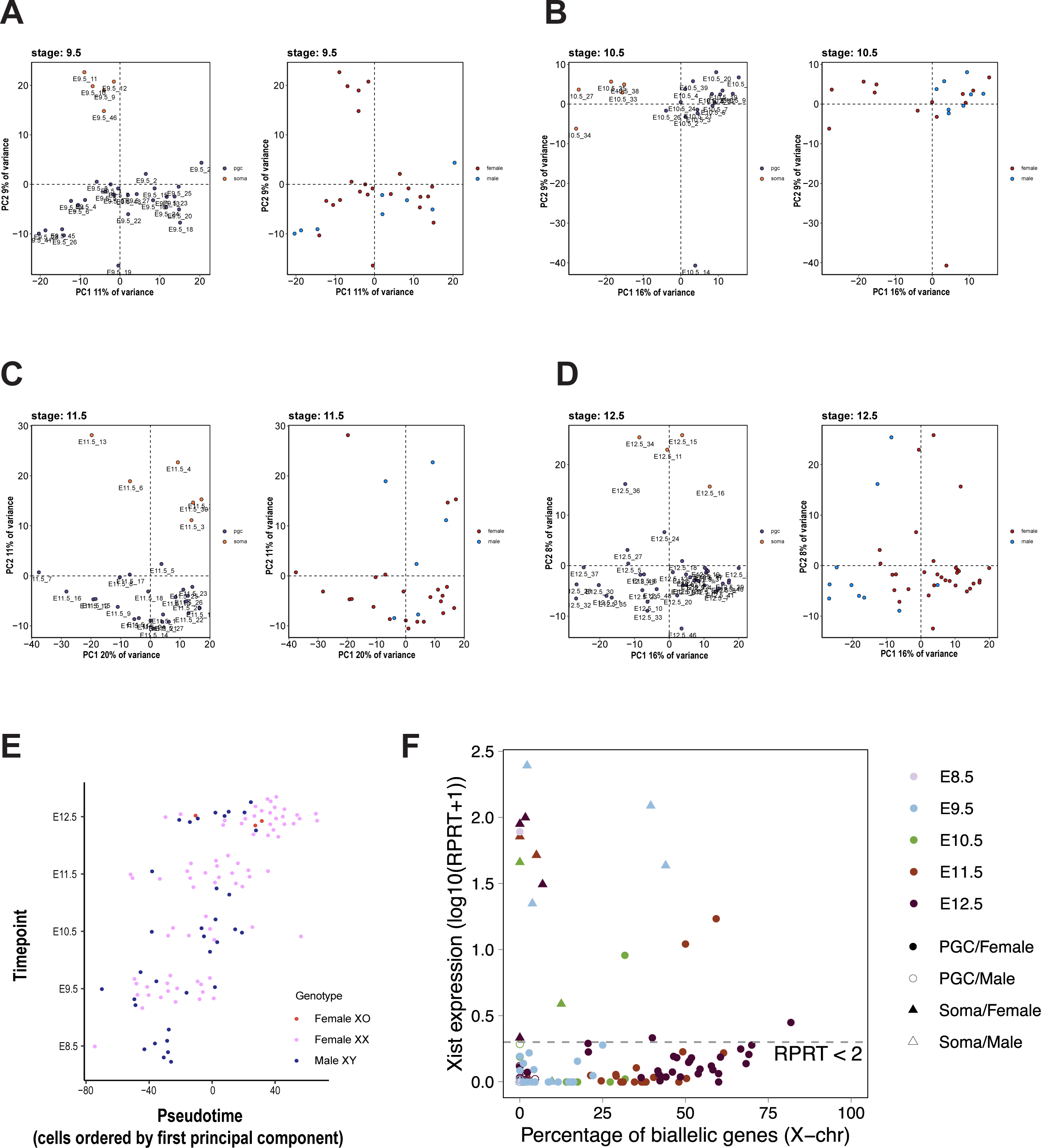
Clustering of PGC by sex. (**A-D**) PCA per developmental stage with sample names, lineage soma versus PGC, and sex information. Cells were clustered by lineage. From E11.5, the cells clustered by sex. **(E)** Pseudotime representation of the scRNA-seq data based on the first principal component for XX females (pink), XO females (red), and XY males (blue). **(F)** Level of *Xist* expression and degree of reactivation in each single cell. Each dot represents a single cell. Most female soma exhibit high *Xist* expression and a low number of biallelically expressed genes. Genes with RPRT expression < 2 were considered to be unexpressed.

**Extended Figure 3.**
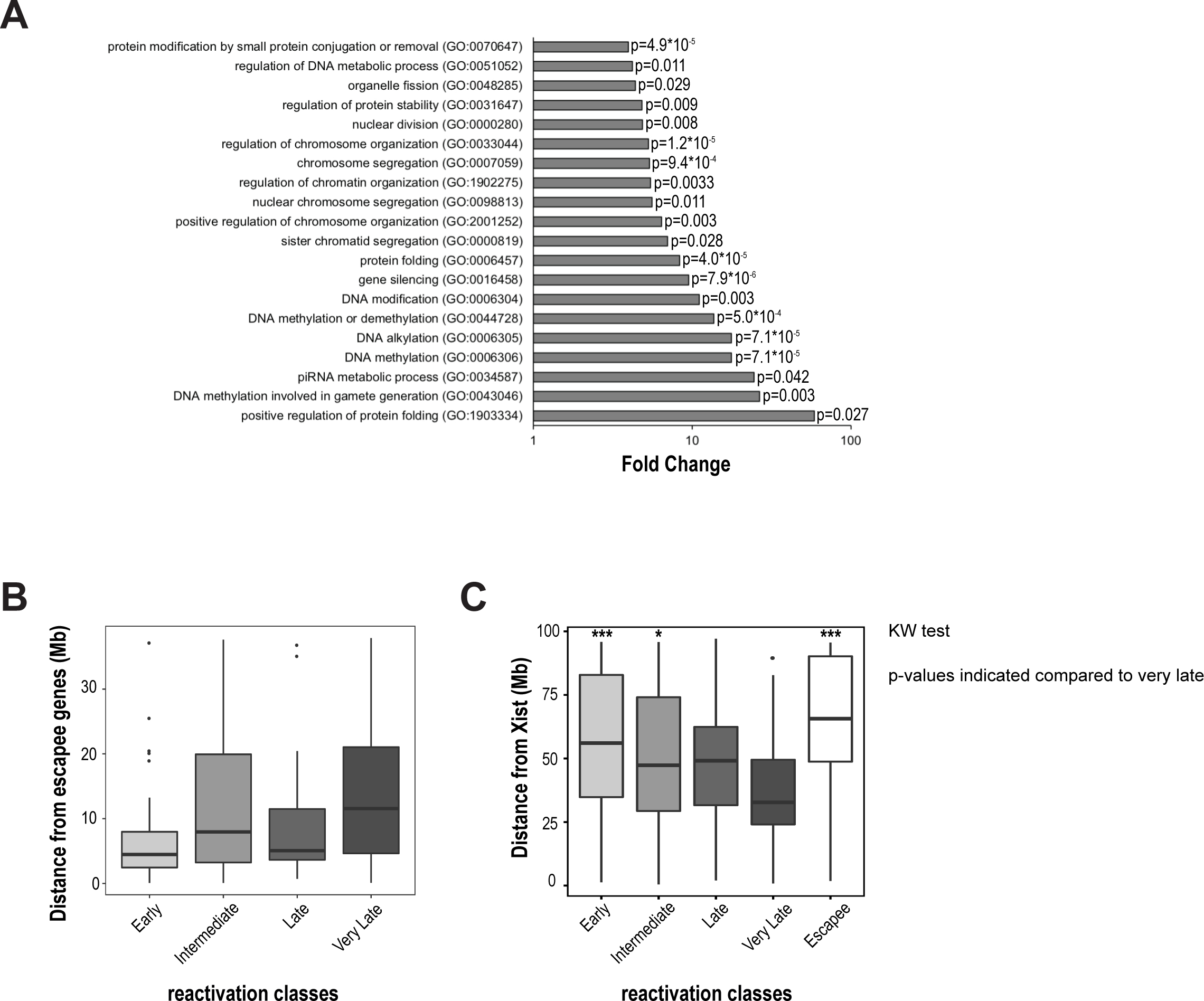
Contribution to kinetics of X-chromosome reactivation. **(A)** Representation of the Gene ontology analysis of Biological process performed on the best correlated genes with X-linked gene reactivation (p-value <0.001). Correlation and anti-correlation between gene expression levels (autosomes and X chromosomes) and the percentage of X-linked gene reactivation (allelic ratio >0.2 for X-linked genes) were measured using Pearson’s correlation and the Benjamini–Hochberg correction. The 20 best enrichment classes (based on fold enrichment) were represented by their p-values. **(B)** Distance to escapee loci. Distribution of genomic distances to escapees (Mb) for different X-linked gene reactivation classes. Transcription Start Site (TSS) of each gene was used to measure the distance from the closest escaping gene. Non-significant by Kruskal-Wallis test. Boxplots represent the medians with lower and upper quartiles. (**C)** Distance to *Xist* genomic locus. Distribution of genomic distances to the *Xist* locus (Mb) for different X-linked gene reactivation classes. The Transcription Start Site (TSS) of each gene was used to measure the distance from the *Xist* locus. Very late reactivated genes were significantly closer to the *Xist* locus than early reactivated (p=0.0041), intermediate-reactivated (p=0.0201), and escapee (p=0.0045) genes, according to the Kruskal-Wallis test. Boxplots represent the medians with lower and upper quartiles.

**Extended Figure 4:**
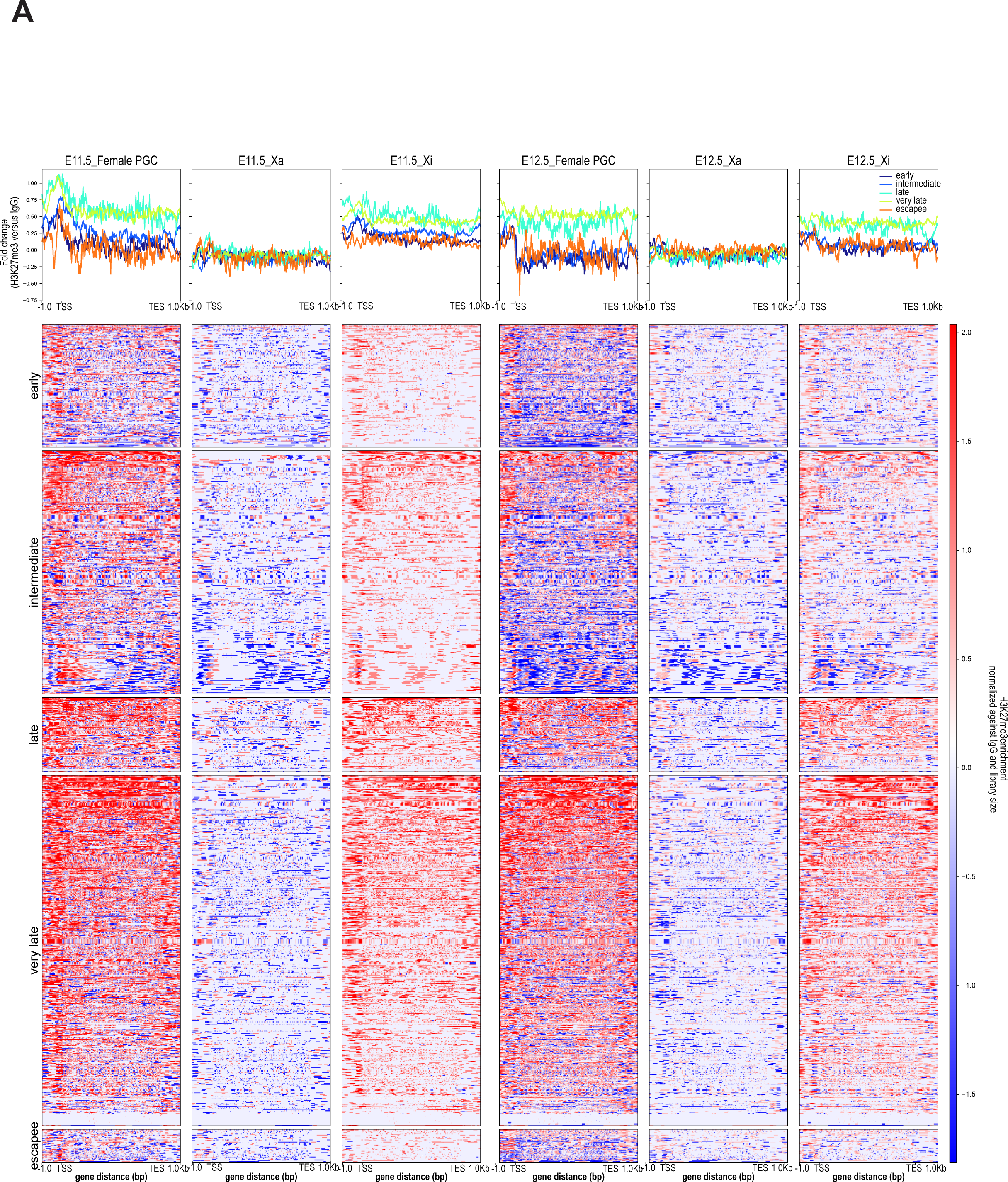
H3K27me3 enrichment by low-input CUT&RUN in E11.5 and E12.5 female PGC. Profile plots and their corresponding heat maps around the transcription start sites (TSS) and transcription end sites (TES) of X-linked genes for H3K27me3 repressive histone marks. Enrichment was extracted for the E11.5 and E12.5 CUT&RUN experiments within a region spanning ± 1 kb around TSS and TES. The blue-to-red gradient indicates low to high enrichment in the corresponding regions ranked by reactivation classes.

**Supplementary Table 1:** Summary of sequenced data including information of scRNA-seq and CUT&RUN datasets.

## References

1. Leitch, H. G., Tang, W. W. C. & Surani, M. A. Primordial Germ-Cell Development and Epigenetic Reprogramming in Mammals. in Current Topics in Developmental Biology vol. 104 149–187 (Elsevier, 2013).

2. Tang, W. W. C., Kobayashi, T., Irie, N., Dietmann, S. & Surani, M. A. Specification and epigenetic programming of the human germ line. Nature Reviews Genetics 17, 585–600 (2016).

3. Kurimoto, K. & Saitou, M. Germ cell reprogramming. Current Topics in Developmental Biology vol. 135 (Elsevier Inc., 2019).

4. Kurimoto, K. & Saitou, M. Epigenome regulation during germ cell specification and development from pluripotent stem cells. Current Opinion in Genetics & Development 52, 57–64 (2018).

5. Hajkova, P. et al. Chromatin dynamics during epigenetic reprogramming in the mouse germ line. Nature 452, 877–81 (2008).

6. Sugimoto, M. & Abe, K. X chromosome reactivation initiates in nascent primordial germ cells in mice. PLoS genetics 3, e116 (2007).

7. Chuva de Sousa Lopes, S. M., et al. X chromosome activity in mouse XX primordial germ cells. PLoS genetics 4, e30 (2008).

8. Severino, J. et al. Controlled X-chromosome dynamics defines meiotic potential of female mouse in vitro germ cells. The EMBO Journal 41, e109457 (2022).

9. Mallol, A., Guirola, M. & Payer, B. PRDM14 controls X - chromosomal and global epigenetic reprogramming of H3K27me3 in migrating mouse primordial germ cells. Epigenetics & Chromatin 12, 1–10 (2019).

10. Sangrithi, M. N. et al. Non-Canonical and Sexually Dimorphic X Dosage Compensation States in the Mouse and Human Germline. Developmental Cell 40, 1–13 (2017).

11. Naik, H. C. et al. Lineage-specific dynamics of erasure of X-upregulation during inactive-X reactivation. 2020.12.23.424181 Preprint at https://doi.org/10.1101/2020.12.23.424181 (2023).

12. Heard, E. & Turner, J. Function of the Sex Chromosomes in Mammalian Fertility. Cold Spring Harb Perspect Biol 1–18 (2011).

13. Lyon, M. F. Gene action in the X-chromosome of the mouse (Mus musculus L.). Nature 190, 372– 3 (1961).

14. Loda, A., Collombet, S. & Heard, E. Gene regulation in time and space during X-chromosome inactivation. Nat Rev Mol Cell Biol 23, 231–249 (2022).

15. Namekawa, S. H., Payer, B., Huynh, K. D., Jaenisch, R. & Lee, J. T. Two-step imprinted X inactivation: repeat versus genic silencing in the mouse. Molecular and cellular biology 30, 3187– 3205 (2010).

16. Borensztein, M. et al. Xist-dependent imprinted X inactivation and the early developmental consequences of its failure. Nature structural & molecular biology 24, 226–233 (2017).

17. Marahrens, Y., Panning, B., Dausman, J., Strauss, W. & Jaenisch, R. Xist-deficient mice are defective in dosage compensation but not spermatogenesis. Genes & Development 11, 156–166 (1997).

18. Engreitz, J. M. et al. The Xist lncRNA exploits three-dimensional genome architecture to spread across the X chromosome. Science (New York, N.Y.) 341, 1237973 (2013).

19. Pandya-Jones, A. et al. A protein assembly mediates Xist localization and gene silencing. Nature (2020) doi:10.1038/s41586-020-2703-0.

20. Strehle, M. & Guttman, M. Xist drives spatial compartmentalization of DNA and protein to orchestrate initiation and maintenance of X inactivation. Curr Opin Cell Biol 64, 139–147 (2020).

21. Chaumeil, J., Baccon, P. L., Wutz, A. & Heard, E. A novel role for Xist RNA in the formation of a repressive nuclear compartment into which genes are recruited when silenced. Genes Dev. 20, 2223– 2237 (2006).

22. Chow, J. C. et al. LINE-1 activity in facultative heterochromatin formation during X chromosome inactivation. Cell 141, 956–69 (2010).

23. Zylicz, J. J. et al. The Implication of Early Chromatin Changes in X Chromosome Inactivation. Cell 176, 182–197.e23 (2019).

24. Monk, M. Methylation and the X chromosome. Bioessays 4, 204–208 (1986).

25. Cheng, S. et al. Single-Cell RNA-Seq Reveals Cellular Heterogeneity of Pluripotency Transition and X Chromosome Dynamics during Early Mouse Development. Cell Reports 26, 2593–2607 (2019).

26. Marks, H. et al. Dynamics of gene silencing during X inactivation using allele-specific RNA-seq. Genome biology 16, 149 (2015).

27. Sousa, L. B. de A. e et al. Kinetics of Xist-induced gene silencing can be predicted from combinations of epigenetic and genomic features. Genome research (2019) doi: 10.1101/gr.245027.118.

28. Mak, W. et al. Reactivation of the paternal X chromosome in early mouse embryos. Science (New York, N.Y.) 303, 666–9 (2004).

29. Borensztein, M. et al. Contribution of epigenetic landscapes and transcription factors to X-chromosome reactivation in the inner cell mass. Nature Communications 8, 1–14 (2017).

30. Pasque, V. et al. X chromosome reactivation dynamics reveal stages of reprogramming to pluripotency. Cell 159, 1681–1697 (2014).

31. Janiszewski, A. et al. Dynamic reversal of random X-Chromosome inactivation during iPSC reprogramming. Genome research 29, 1659–1672 (2019).

32. de Napoles, M., Nesterova, T. & Brockdorff, N. Early loss of Xist RNA expression and inactive X chromosome associated chromatin modification in developing primordial germ cells. PLoS One 2, e860 (2007).

33. Mallol, A., Guirola, M. & Payer, B. PRDM14 controls X - chromosomal and global epigenetic reprogramming of H3K27me3 in migrating mouse primordial germ cells. Epigenetics & Chromatin 12, 1–10 (2019).

34. Navarro, P. et al. Molecular coupling of Xist regulation and pluripotency. Science (New York, N.Y.) 321, 1693–5 (2008).

35. Payer, B. pdf et al. Tsix RNA and the Germline Factor, PRDM14, Link X Reactivation and Stem Cell Reprogramming. Molecular Cell 52, 1–14 (2013).

36. Wang, M., Lin, F., Xing, K. & Liu, L. Random X-chromosome inactivation dynamics in vivo by single-cell RNA sequencing. BMC Genomics 18, 90 (2017).

37. Keane, T. M. et al. Mouse genomic variation and its effect on phenotypes and gene regulation. Nature 477, 289–94 (2011).

38. Frazer, K. A. et al. A sequence-based variation map of 8. 27 million SNPs in inbred mouse strains. Nature >448, 1050–1053 (2007).

39. Payer, B. et al. Generation of stella-GFP Transgenic Mice : A Novel Tool to Study Germ Cell Development. Genesis 44, 75–83 (2006).

40. Yeom, Y. I. et al. Germline regulatory element of Oct-4 specific for the totipotent cycle of embryonal cells. Development 122, 881–94 (1996).

41. Tang, F. et al. mRNA-Seq whole-transcriptome analysis of a single cell. Nature methods 6, (2009).

42. Borensztein, M., Syx, L., Servant, N. & Heard, E. Transcriptome Profiling of Single Mouse Oocytes. Methods in Molecular Biology 1818, 51–65 (2018).

43. Mayère, C. et al. Single-cell transcriptomics reveal temporal dynamics of critical regulators of germ cell fate during mouse sex determination. The FASEB Journal 35, e21452 (2021).

44. Yamaji, M. et al. DND1 maintains germline stem cells via recruitment of the CCR4-NOT complex to target mRNAs. Nature 543, 568–572 (2017).

45. Bertozzi, T. M., Elmer, J. L., Macfarlan, T. S. & Ferguson-Smith, A. C. KRAB zinc finger protein diversification drives mammalian interindividual methylation variability. Proc. Natl. Acad. Sci. U.S.A. 117, 31290–31300 (2020).

46. Dehghanian, F., Bovio, P. P., Hojati, Z. & Vogel, T. ZFP982 confers mouse embryonic stem cell characteristics by regulating expression of Nanog, Zfp42 and Dppa3. 2020.06.03.131847 Preprint at https://doi.org/10.1101/2020.06.03.131847 (2020).

47. Yu, L. et al. Loss of ESRP1 blocks mouse oocyte development and leads to female infertility. Development 148, dev196931 (2021).

48. Zhao, Z.-H. et al. Single cell RNA sequencing reveals the landscape of early female germ cell development. FASEB Journal 2020.05.09.085845 (2020) doi:10.1101/2020.05.09.085845.

49. Cattanach, B. M. & Williams, C. E. Evidence of non-random X chromosome activity in the mouse. Genetics Research 19, 229–240 (1972).

50. Calaway, J. D. et al. Genetic Architecture of Skewed X Inactivation in the Laboratory Mouse. PLoS Genet 9, e1003853 (2013).

51. Berletch, J. B. et al. Escape from X Inactivation Varies in Mouse Tissues. PLOS Genetics 11, e1005079 (2015).

52. Pandya-Jones, A. & Plath, K. The lnc between 3D chromatin structure and X chromosome inactivation. Seminars in Cell and Developmental Biology 56, 35–47 (2016).

53. Kobayashi, H. et al. High-resolution DNA methylome analysis of primordial germ cells identifies gender-specific reprogramming in mice. Genome Res. 23, 616–627 (2013).

54. Hill, P. W. S. et al. Epigenetic reprogramming enables the primordial germ cell-to-gonocyte transition Europe PMC Funders Group. Nature 555, 392–396 (2018).

55. Richart, L. et al. XIST loss impairs mammary stem cell differentiation and increases tumorigenicity through Mediator hyperactivation. Cell 185, 2164–2183.e25 (2022).

56. Generoso, S. F. et al. Cohesin controls X chromosome structure remodeling and X-reactivation during mouse iPSC-reprogramming. Proceedings of the National Academy of Sciences 120, e2213810120 (2023).

57. Camlin, N. J., McLaughlin, E. A. & Holt, J. E. Kif4 Is Essential for Mouse Oocyte Meiosis. PLoS One 12, e0170650 (2017).

58. Luoh, S.-W. et al. Zfx mutation results in small animal size and reduced germ cell number in male and female mice. Development 124, 2275–2284 (1997).

59. Lan, Z. J., Xu, X. & Cooney, A. J. Differential oocyte-specific expression of Cre recombinase activity in GDF-9-iCre, Zp3cre, and Msx2Cre transgenic mice. Biol Reprod 71, 1469–1474 (2004).

60. Szabo, P. E., Hübner, K., Schöler, H. & Mann, J. R. Allele-specific expression of imprinted genes in mouse migratory primordial germ cells. Mechanisms of development 115, 157–160 (2002).

61. Tang, F., Lao, K. & Surani, M. A. Development and applications of single-cell transcriptome analysis. Nature methods 8, (2011).

62. Skene, P. J. & Henikoff, S. An efficient targeted nuclease strategy for high-resolution mapping of DNA binding sites. eLife 6, 1–35 (2017).

63. Prakash, S. A. & Barau, J. Chromatin Profiling in Mouse Embryonic Germ CellsEmbryonic germ cells by CUT&RUN. in Epigenetic Reprogramming During Mouse Embryogenesis: Methods and Protocols (eds. Ancelin, K. & Borensztein, M.) 253–264 (Springer US, 2021). doi:10.1007/978-1-0716-0958-3_17.

64. Dura, M. et al. DNMT3A-dependent DNA methylation is required for spermatogonial stem cells to commit to spermatogenesis. Nat Genet 54, 469–480 (2022).

65. Ewels, P., Magnusson, M., Lundin, S. & Käller, M. MultiQC: summarize analysis results for multiple tools and samples in a single report. Bioinformatics 32, 3047–3048 (2016).

66. Martin, M. Cutadapt removes adapter sequences from high-throughput sequencing reads. EMBnet.journal 17, 10–12 (2011).

67. Shen, W., Le, S., Li, Y. & Hu, F. SeqKit: A Cross-Platform and Ultrafast Toolkit for FASTA/Q File Manipulation. PLOS ONE 11, e0163962 (2016).

68. Krueger, F., James, F., Ewels, P., Afyounian, E. & Schuster-Boeckler, B. FelixKrueger/TrimGalore: v0.6.7 - DOI via Zenodo. (Zenodo, 2021). doi:10.5281/zenodo.5127899.

69. Langmead, B. & Salzberg, S. L. Fast gapped-read alignment with Bowtie 2. Nat Methods 9, 357– 359 (2012).

70. Krueger, F. & Andrews, S. R. SNPsplit: Allele-specific splitting of alignments between genomes with known SNP genotypes. F1000Res 5, 1479 (2016).

71. Danecek, P. et al. Twelve years of SAMtools and BCFtools. Gigascience 10, giab008 (2021).

72. Ramírez, F. et al. deepTools2: a next generation web server for deep-sequencing data analysis. Nucleic Acids Res 44, W160–W165 (2016).

73. Zhang, Y. et al. Model-based Analysis of ChIP-Seq (MACS). Genome Biology 9, R137 (2008).

74. Liao, Y., Smyth, G. K. & Shi, W. featureCounts: an efficient general purpose program for assigning sequence reads to genomic features. Bioinformatics 30, 923–930 (2014).

75. Smallwood, S. a et al. Single-cell genome-wide bisulfite sequencing for assessing epigenetic heterogeneity. Nature Methods 11, 817–20 (2014).

76. Walter, M., Teissandier, A., Pérez-Palacios, R. & Bourc’his, D. An epigenetic switch ensures transposon repression upon dynamic loss of DNA methylation in embryonic stem cells. eLife 5, e11418 (2016).

77. Krueger, F. & Andrews, S. R. Bismark: a flexible aligner and methylation caller for Bisulfite-Seq applications. Bioinformatics 27, 1571–1572 (2011).

